# BAP1 dysregulation impairs trophoblast differentiation and contributes to placental dysfunction in preeclampsia

**DOI:** 10.1101/2025.11.28.691117

**Authors:** Paula Doria-Borrell, Ana Ferrero-Micó, Sergio Navarro-Serna, Maravillas Mellado-López, Johanna Grinat, Ciara N Murphy, Lina Youssef, Fàtima Crispi, Tu’uhevaha J Kaitu’u-Lino, Vicente Pérez-García

## Abstract

Preeclampsia, a life-threatening hypertensive disorder of pregnancy, is a leading cause of maternal and perinatal morbidity and mortality. Its early-onset form (EO-PE), requiring delivery before 34 weeks of gestation, is particularly severe and closely linked to defective trophoblast differentiation. Here, we identify BRCA1-associated protein 1 (BAP1) and its cofactors ASXL2 and ASXL3 are upregulated in EO-PE placentas. Enforced BAP1 expression in human trophoblast stem cells reinforced epithelial identity, enhanced adhesion, and impaired both extravillous trophoblast differentiation and syncytiotrophoblast formation. Integrated transcriptomic and proteomic analyses revealed suppression of lineage-specific pathways alongside maintenance of progenitor-like and pro-inflammatory signatures. In trophoblast organoids, excess of BAP1 disrupted syncytial maturation and induced interferon-driven pathways overlapping with EO-PE transcriptomes. Together, these findings establish BAP1 as a key regulator of human trophoblast differentiation and implicate its dysregulation in the pathogenesis of EO-PE, providing mechanistic insight into the cellular basis of placental dysfunction.

## Introduction

The human placenta is a transient yet essential organ that supports fetal development by mediating maternal–fetal exchange and regulating immune and endocrine interactions throughout gestation. Its architecture and function are established early in pregnancy through the expansion and differentiation of cytotrophoblasts (CTBs), which give rise to syncytiotrophoblasts (STBs) and extravillous trophoblasts (EVTs). STBs form the outer multinucleated layer of the chorionic villi, mediating nutrient and gas exchange, while EVTs migrate into the maternal decidua to remodel uterine spiral arteries and ensure adequate blood flow to the developing fetus. The differentiation of CTBs into these specialized trophoblast lineages is therefore fundamental for normal placental development and function^1–3^. Early studies using isolated primary CTBs and first-trimester placental explants yielded valuable insights into human trophoblast biology, including the processes of syncytial fusion and EVT invasion. Although informative, these models are restricted by limited tissue availability, intrinsic cellular heterogeneity, and their inability to sustain proliferative trophoblast populations in culture. As a result, they provide only a narrow window into how trophoblast progenitor states are defined, stabilized, or perturbed during early placental development^4^. A major advance came with the derivation of human trophoblast stem cells (hTSCs) from blastocysts and first-trimester villi, which established a renewable source of trophoblast progenitors capable of differentiating into both major placental lineages^5^. The subsequent development of trophoblast organoids from first-trimester tissue further addressed the need for physiologically relevant in vitro systems^6,7^. These three-dimensional structures recreate core architectural and functional hallmarks of the trophoblast epithelium, including lineage diversification, spatial organization, and robust endocrine activity.

Together, hTSC and trophoblast organoid platforms have transformed experimental access to human-specific mechanisms governing trophoblast fate. Their genetic tractability and lineage fidelity now allow systematic exploration of how molecular regulators control the balance between self-renewal, differentiation, and programmed cell death. These models therefore constitute an essential framework for interpreting how dysregulated trophoblast programs contribute to placental dysfunction and pregnancy disorders.

Defective trophoblast differentiation is a hallmark of placental disorders, most notably preeclampsia (PE), a hypertensive pregnancy complication with major maternal and perinatal morbidity^8,9^. PE is increasingly recognized as a heterogeneous disorder with distinct early- and late-onset subtypes^10^. Early-onset PE (EO-PE) occurs before 34 weeks of gestation and is primarily driven by defective placentation, including shallow EVT invasion, impaired spiral artery remodeling, and diminished syncytialization^11–13^. EO-PE placentas also exhibit heightened oxidative stress and activation of the unfolded protein response (UPR), which suppresses non-essential protein synthesis in an attempt to restore endoplasmic reticulum (ER) homeostasis^14^. Consistent with this heightened stress, shedding of syncytiotrophoblast-derived micro- and nanovesicles, including exosomes carrying regulatory microRNAs, is markedly increased in EO-PE compared to both healthy pregnancies and late-onset disease^15–18^. In contrast, late-onset PE (LO-PE), which manifests after 34 weeks, is thought to reflect a mismatch between normal placental aging and the increasing metabolic demands of the fetus in the context of maternal predisposition to cardiovascular or metabolic disease^10^. LO-PE is frequently associated with maternal inflammation, elevated BMI, and higher arterial pressure, and although placental dysfunction is present, it is generally less severe than in EO-PE^19^. Despite these advances, the regulators that govern trophoblast lineage decisions in EO-PE, and differentiate it from the largely maternal-driven pathology of LO-PE, remain unclear.

Our previous work in mice identified BRCA1-associated protein 1 (BAP1) as a key regulator of placentation^20^. BAP1 functions as the catalytic subunit of the Polycomb Repressive-DUB (PR-DUB) complex by interacting with additional sex comb-like (ASXL1/2/3) proteins to deubiquitinate histone H2A at lysine 119 (H2AK119), maintaining its spatial distribution and exerting its tumour suppressor activity^21–23^. Mouse models deficient in Bap1 exhibit severe placental abnormalities and midgestational lethality^20^. In both murine and human placentas, BAP1 is highly expressed in proliferative CTBs and downregulated during differentiation into EVTs and STBs, suggesting a conserved role in lineage regulation^24^.

Here, we dissect the role of BAP1 in human trophoblast development using CRISPR/Cas9 knockout technology and genetic overexpression in hTSCs. We show that BAP1 is upregulated in EO-PE placentas, supporting its pathological relevance. We demonstrate that high BAP1 expression sustains epithelial features, impedes differentiation into both EVTs and STBs, and mimics molecular signatures of EO-PE. Our findings highlight proper BAP1 modulation as a critical step in trophoblast differentiation, with direct implications for the etiology of preeclampsia.

## MATERIAL AND METHODS

### Human Samples

#### Melbourne Cohort

Details on the number and characteristics of the patients and control subjects are provided in Supplementary File 7. The study was approved by the Mercy Health Human Research Ethics Committee (R11/34) and written informed consent was obtained from all participants prior to sample collection Placental tissue samples were collected in Melbourne within 30 minutes of delivery and snap-frozen at −80 °C for subsequent RNA extraction. Total mRNA was extracted and reverse-transcribed into cDNA using the High-Capacity cDNA Reverse Transcription Kit (Applied Biosystems, Life Technologies, Carlsbad, CA) with 400 ng of input mRNA, according to the manufacturer’s instructions. The thermal cycling conditions were: 25 °C for 10 minutes, 37 °C for 60 minutes, and 85 °C for 5 minutes. cDNA was stored at −20 °C. Quantitative PCR was performed using a CFX384 real-time PCR system (Bio-Rad) and Fast SYBR Master Mix (Thermo Fisher Scientific, Scoresby, Australia). The cycling conditions were: 95 °C for 20 seconds, followed by 40 cycles of 95 °C for 1 second and 60 °C for 20 seconds, with a melt curve from 65 °C to 95 °C in 0.5 °C increments every 5 seconds.

### Barcelona Cohort

Eighteen singleton pregnancies were prospectively recruited at the Department of Maternal-Fetal Medicine, Hospital Clinic (Barcelona, Spain) between 2017 and 2021. The cohort included normotensive pregnancies (n = 9) and cases of preeclampsia (n = 9), defined as systolic blood pressure >140 mmHg and/or diastolic blood pressure >90 mmHg on two separate occasions (≥4 hours apart) after 20 weeks of gestation, accompanied by proteinuria (>300 mg/24 h or protein/creatinine ratio >300 mg/g) or signs of end-organ dysfunction^25^. Early-onset preeclampsia was defined as requiring delivery before 34 weeks’ gestation; late-onset cases delivered at or after 34 weeks. Controls were matched for baseline characteristics, including several preterm deliveries due to spontaneous labor or non-hypertensive indications. Exclusion criteria included fetal anomalies, chromosomal abnormalities, and intrauterine infections. Gestational age was determined by first-trimester crown-rump length. Ethics approval was granted by the Hospital Clinic Ethics Committee (HCB-2016-0830), and written informed consent was obtained from all participants.

Placental tissues were collected at delivery, snap-frozen, and stored at −80 °C. Samples were handled on dry ice. Tissue exceeding ∼30 mg was trimmed on a pre-chilled P60 plate and transferred to matrix A-tubes using a 21G needle. Samples were homogenized using the FastPrep-24 5G system (MP Biomedicals) in 300 μL of TRIzol or Triplex + CI. Four homogenization cycles were performed, with the rotor placed on ice between runs. Samples were centrifuged at 14,000 g for 10 minutes at 4 °C. The supernatant was transferred to fresh tubes. For viscous samples, 200 μL of additional TRIzol was added, followed by vortexing and repeat centrifugation under identical conditions. Final supernatants were stored at −80 °C.

For protein extraction, tissues were homogenized in RIPA buffer (1X) supplemented with protease/phosphatase inhibitors (CI) and PMSF. Homogenization was conducted using the same FastPrep-24 protocol as above. After centrifugation (14,000 g, 10 minutes, 4 °C), the supernatant was collected, sonicated for 5 minutes, and quantified. Outlier samples (*n*=4) were diluted with an additional 150 μL RIPA + CI + PMSF before re-quantification. Proteins were resolved on 7% SDS-PAGE gels and transferred at 290 mA for 90 minutes. Standard western blot procedures were followed for membrane blocking and antibody incubation.

An alternative extraction method using an Ultraturrax homogenizer (Polytron) was employed for optimization. Placental fragments were homogenized in 350 μL of RIPA + CI + PMSF on ice with two 20-second pulses separated by a 30-second cooling interval. The homogenate was centrifuged (maximum speed, 10 minutes, 4 °C), and the supernatant was processed for protein quantification and western blotting as described above.

### Culture and differentiation of human trophoblast cell lines

BTS11 blastocyst-derived hTSCs were obtained from Professor Takahiro Arima and cultured following the protocol described by Okae et al. (2018), with an increased concentration of CHIR99021^5^.In summary, hTSCs were plated at a density of 0.5–1×10⁶ cells per well on six-well plates coated with 5 mg/mL collagen IV (Cultrex 3410-010-02). The cells were maintained in 2 mL of TS medium, which consisted of DMEM/F12 (Gibco 31331-028) supplemented with 0.3% BSA (Sigma A8412), 1% ITS-X supplement (Gibco 51500-056), 0.1 mM 2-mercaptoethanol (Gibco 31350), 0.2% FBS (Gibco 10270), 100 µg/mL Primocin (Invivogen ant-pm-1), 1.5 µg/mL L-ascorbic acid (Sigma A8960), 50 ng/mL EGF (Peprotech AF-100-15), 4 µM CHIR99021 (Tocris 4423), 0.5 µM A83-01 (Systems Biosciences ZRD-A8-02), 1 µM SB431542 (Tocris 1614), 0.8 mM VPA (Sigma P4543), and 5 µM Rock Inhibitor Y27632 (Millipore 688000). Cultures were maintained at 37°C with 5% CO₂, and the medium was replaced every two days. For passaging, cells were detached using TrypLE (Gibco 12604-013) for 10–15 minutes at 37°C.

For differentiation into extravillous trophoblasts (EVTs), the method followed was based on the protocol described in^24^. Briefly, hTSCs at 80% confluence were detached using TrypLE and resuspended as single cells. A total of 0.75×10⁵ cells were plated onto six-well plates coated with 1 µg/mL collagen IV and cultured in 2 mL of EVT medium. This medium consisted of DMEM/F12 supplemented with 0.1 mM 2-mercaptoethanol, 0.5% penicillin-streptomycin, 0.3% BSA, 1% ITS-X supplement, 100 ng/mL NRG1, 7.5 µM A83-01, 2.5 µM Y27632, and 4% KSR. Additionally, Matrigel was included at a final concentration of 2%. On day 3, the medium was changed to EVT medium without NRG1, with Matrigel adjusted to 0.5%. On day 6, the medium was refreshed again, this time omitting both NRG1 and KSR while keeping Matrigel at 0.5%. The differentiation process was assessed on day 8.

To induce syncytiotrophoblast (STB) differentiation, hTSCs at approximately 80% confluence were dissociated into single cells using TrypLE. A total of 1.5×10⁵ cells were then seeded onto a fresh six-well plate pre-coated with 3 µg/mL collagen IV. The cells were cultured in STB differentiation medium, as outlined by Okae et al. (2018), which included DMEM/F12 GlutaMAX (Gibco 31331-028) with 0.1 mM 2-mercaptoethanol (Gibco 31350), 4% KSR, 0.3% BSA (Sigma A8412), 1% ITS-X supplement (Gibco 51500-056), 2.5 µM Y27632 (Millipore 688000), 2 µM forskolin (Sigma 93049), and 0.5% Primocin (Invivogen ant-pm-1). The medium was refreshed on day 2, and analyses were performed on day 4.

### Organoid culture

The transition from 2D cultures to 3D trophoblast organoids was carried out by seeding a single-cell suspension onto 25 µL Matrigel droplets in 48-well plates. The culturing process followed the established protocol described in^26^. After incubating the Matrigel droplets with organoids for 10 minutes at 37°C, cultures were overlaid with trophoblast organoid medium (TOM). This medium comprised Advanced DMEM/F12 (Thermo Fisher Scientific), N2 supplement (Thermo Fisher Scientific, 17502048), B27 supplement without vitamin A (Thermo Fisher Scientific, 12587010; used as per the manufacturer’s instructions), 100 µg/mL Primocin (InvivoGen, ant-pm), 1.25 mM N-acetyl-L-cysteine (Sigma-Aldrich, A9165), 2 mM L-glutamine (Sigma-Aldrich, 25030024), 50 ng/mL recombinant human EGF (Peprotech, AF10015), 1.5 µM CHIR99021 (Tocris, 4423), 80 ng/mL recombinant human R-spondin-1 (R&D Systems, 4645-RS-01M/CF), 100 ng/mL recombinant human FGF-2 (Peprotech, 100-18B), 50 ng/mL recombinant human HGF (Peprotech, 100-39), 500 nM A83-01 (Tocris, 2939), 2.5 µM prostaglandin E2 (Sigma-Aldrich, P0409), and 2 µM Y-27632 (Millipore, 688000).

To passage the organoids, Matrigel droplets were collected into 1.5 mL tubes and mechanically dissociated through repeated pipetting (∼400 times) until the organoids were broken down. The resulting organoid suspension was then washed with Advanced DMEM/F12 and reseeded into fresh Matrigel droplets. For pellet collection, organoid droplets were transferred into 1.5 mL tubes containing Cell Recovery Solution and incubated for 30 minutes at 4°C. After washing with PBS, organoids were either analyzed immediately or stored at -80°C for later use.

### CRISPR-Cas9 mediated knockdown in hTSCs

For CRISPR/Cas9-mediated knockdown of BAP1, sgRNA oligonucleotides (Integrated DNA Technologies) were designed to target regions upstream and downstream of BAP1 exon 4 using http://crispor.tefor.net/, as described in Doria-Borrell et al. (2024). A total of four sgRNAs were designed: two upstream (1A and 1B) and two downstream (2A and 2B) of exon 4. The sgRNA sequences were as follows: 1A, 5′-TTTGCACTGCGTCATCACTC-3′; 1B, 5′-TGAGTGATGACGCAGTGCAA-3′; 2A, 5′-CAGAGCAGGTCAGTCATATC-3′; and 2B, 5′-AGTTCAGTTCGTTCTGCCAG-3′.

Each sgRNA sequence was cloned into either pSpCas9(BB)-2A-GFP (PX458; Addgene plasmid #48138) or pSpCas9(BB)-2A-Puro V2.0 (PX459; Addgene plasmid #62988). Human trophoblast stem cells (hTSCs) were transfected with plasmid combinations of (1A–puromycin + 2A–GFP) or (1A–puromycin + 2B–GFP) using Opti-MEM (Thermo Fisher Scientific) and Lipofectamine 2000 reagent according to the manufacturer’s instructions (Thermo Fisher, 11668019). Forty-eight hours after transfection, GFP⁺ cells were sorted using a FACSAria II system and replated onto 6-well plates according to cell number. Then, cells were treated with 1 µg/mL puromycin for 3 days.

### Lentivirus over-expression in hTSCs

BTS11 cells were transduced with lentiviral particles of pLV[Exp]-EGFP:T2A:Puro-CMV>ORF_Empty (Control) or (pLV [Exp]-Puro- CMV > EGFP: hBAP1[NM_004656.4] or pLV[Exp]-EGFP/Puro-EF1A>hBAP1[NM_004656.4] to overexpress BAP1. In both cases, the lentiviral particles were packaged by VectorBuilder. Cells were transduced with the lentivirus supplemented with 8ug/mL polybrene (Millipore, TR-1003-G) for 24 hours. After 48 hours, cells were selected by sorting and collecting all GFP+ cells in one P48 cell plate. After cell expansion, we selected again by adding 1ug/mL puromycin to the medium for 7 days.

### Proliferation assay

Trophoblast cells (150.000 cells/well) were seeded in a 6-well plate pre-coated with 5 mg/mL Col IV (Cultrex 3410-010-02). Cells were collected every 24 hours using TrypLE (Gibco 12604-013) for 10 to 15 min at 37 °C and viable cells were counted using trypan blue exclusion on a LUNA II Automated Cell Counter (Logos Biosystems) according to the manufacturer’s instructions. Data was analysed using two-way ANOVA with Sidák’s multiple comparisons test (p-value < 0.05) using GraphPad Prism 9.

### Cell adhesion assay

The adhesion capacity of control, BAP1 knockdown (BAP1-KD), and BAP1-overexpressing (BAP1-OE) human trophoblast stem cells (hTSCs) was assessed using the Vybrant Cell Adhesion Assay Kit (Thermo Fisher Scientific, V13181), following the manufacturer’s instructions and as previously described^27^.

Briefly, a 96-well tissue culture plate was coated with 5 mg/mL collagen IV (Cultrex, 3410-010-02). Cells were resuspended in DMEM/F12 GlutaMAX (Gibco, 31331-028) containing 5 μL of Vybrant labeling reagent and incubated for 30 minutes at 37 °C. Labeled cells were washed with phosphate-buffered saline (PBS; Cytiva, SH30028.02) and resuspended in hTSC complete medium.

A total of 5 × 10⁴ labeled cells were seeded per well into four replicate collagen-coated wells and three uncoated wells (negative control) and allowed to adhere for 4 hours at 37 °C. Following incubation, wells were washed three times with PBS to remove non-adherent cells, and fluorescence was measured at 517 nm using a PHERAstar FS plate reader (BMG Labtech). Data were analyzed using a two-tailed Student’s *t*-test, with statistical significance set at *p* < 0.05 (GraphPad Prism v9).

### Gelatin zymography

Matrix metalloproteinase (MMP) activity in hTSCs and differentiated extravillous trophoblasts (EVTs) was assessed by gelatin zymography. hTSCs were cultured to high confluency and EVTs were collected on day 7 of differentiation. At this stage, cells were cultured in serum-free medium for 24 hours. Conditioned media were then collected and total protein concentrations were determined using a calibration curve.

For each sample, 10 μL of conditioned medium was mixed with 5× non-reducing sample buffer (4% SDS, 20% glycerol, 0.01% bromophenol blue, 125 mM Tris-HCl, pH 6.8) and loaded onto SDS–polyacrylamide gels (7.5% separating gel and 4% stacking gel) containing gelatin as a substrate. Electrophoresis was carried out at 100 V until the protein front migrated into the separating gel, followed by a voltage increase to 140 V until completion.

Gels were washed twice for 30 minutes in washing buffer (2.5% Triton X-100, 50 mM Tris-HCl, pH 7.5, 5 mM CaCl₂, 1 μM ZnCl₂) at room temperature with agitation to remove SDS and restore enzyme activity. Gels were then incubated for 5–10 minutes in incubation buffer (1% Triton X-100, 50 mM Tris-HCl, 5 mM CaCl₂, 1 μM ZnCl₂) at 37 °C, followed by a 24-hour incubation in fresh incubation buffer at 37 °C.

Following incubation, gels were stained in 40% methanol, 10% acetic acid, and 0.5 g/L Coomassie Brilliant Blue R-250 for 30–60 minutes at room temperature with agitation, then destained in 40% methanol and 10% acetic acid until clear bands of gelatinolytic activity were visible. Gels were imaged using the ImageQuant system, and MMP activity was quantified by densitometry using ImageJ software (NIH)

### RNA extraction and RT-qPCR

Total RNA was extracted using RNeasy mini-Kit and RNase-free DNase (QIAGEN 74106). First, cDNA was synthesized from total RNA using PrimeScript RT Master Mix (Takara Bio RR036A), and real-time PCR reaction was done with the TB Green Premix Ex Taq (Takara Bio RR420L) on a LC480 II thermocycler in a 384 plate. The geometric medium of *YWHAZ* and *TBP* was used as housekeeping genes. Full list of primers can be consulted in Supplementary File 6. For statistical analysis, a two-way ANOVA with Tukey’s multiple comparisons test (p-value < 0.05) were performed to calculate statistical significance of expression differences using GraphPad Prism 9.

### Western Blot

Samples collected for Western blot analysis were lysed in RIPA buffer (50mM tris-HCl, 150mM NaCl, IGEPAL CA-630, 0.75% sodium deoxycholate, 0.1% SDS), containing a protease inhibitor cocktail (Sigma P2714), and incubated at 4 °C for 1 h, followed by centrifugation (9300 × g, 10 min). Western blotting was performed as in^28^. Blots were probe against the antibodies anti-SDC1 1:1000 (Abcam, ab128936), anti-HSP90 1:5000 (Cell Signaling, 4877S), anti-TUBULIN (1:5000, Abcam, ab6160), anti-BAP1 1:1000 (Cell signalling, 1318S), anti-ASXL1 1:1000 (Abnova, H00171023-M05), anti-ASXL2 1:1000 (Bethyl, A302-037A), anti-ASXL3 1:500 (MyBioSource, MBS5401046), anti-HLA-G 1:1000 (Biorad, MCA2043), anti-ACTIN 1:5000 (Biorad, ab6276), followed by horseradish peroxidise-conjugated secondary antibodies anti-rabbit (Bio-Rad 170–6515) and anti-mouse (Bio-Rad 170–6516, all 1:3000) (Supplementary material 1). Detection was carried out with enhanced chemiluminescence reaction (GE Healthcare RPN2209) on X-ray films. The intensity of the bands was quantified using ImageJ software.

### Immunofluorescence staining

Cells were fixed with 4% paraformaldehyde (PFA) in phosphate-buffered saline (PBS) for 10 min and permeabilized with PBS, 0.3% Triton X-100 for 10 min. Blocking was carried out with PBS, 0.1% Tween 20, 1% BSA (PBT/BSA) for 30 min, followed by antibody incubation for 60 min. Primary antibodies and dilutions (in PBT/BSA) were E-cadherin (CDH1) 1:100 (BD Biosciences, 610181),), CGb 1:100 (Abcam, ab9582), SDC1) 1:100 (Abcam, AB128936), and DESMOPLAKIN 1:100 (Abcam, ab71690), followed by secondary antibodies Alexa Fluor 568 or 647 (Thermo Fisher Scientific) diluted 1:400 in blocking buffer.

Organoid staining was performed following a modification of the protocol previously described^24^. Briefly, matrigel was removed from organoids using Cell Recovery Solution (Corning) and fixed for 1 hr in 4% PFA at RT. After three washes (15 min) with PBS supplemented with 3mg/ml poly-vinylpyrrolidone (Sigma, P0930), organoids were permeabilized with PBS containing 5% DMSO, 0.5% Triton X-100, 0.1% BSA, 0.01% Tween 20 for 3 hr. Then, organoids were blocked overnight at 4°C in permeabilization buffer, containing 2% Fetal Bovine Serum. Organoids were incubated overnight at 4°C with antibodies against CDH1 at 1:100 (BD Biosciences, 610181), CGb at 1:100 (Abcam, ab9582), and SDC1 at 1:100 (Abcam, AB128936), in blocking buffer, followed by three washes in blocking buffer for 1 hr. Then, organoids were incubated overnight with secondary Alexa Fluor 568 or 647 (Thermo Fisher Scientific) antibodies diluted 1:400 in blocking buffer. Lastly, organoids were washed three times for 1 hour in blocking buffer and nuclei were counter-stained with DAPI for 10 min. Organoids were mounted in Vectashield (Vector Laboratories, H-1000), surrounded by spacer drops of vaseline for the coverslip, to immobilize organoids. Images were taken with a LSM900 Vertical confocal microscope.

### Analysis of organoid perimeter

Measurements of total organoid and syncytiotrophoblast perimeters were performed using bright field images acquired with an EVOS microscope. Image analysis was conducted in ImageJ, where the perimeter of each organoid was manually outlined and measured. To ensure accuracy and account for variability, each organoid was measured three times, and the mean perimeter was calculated. Statistical comparison was Student’s two-tailed t-tests, performed using GraphPad Prisms (v9.4.1). Statistical significance was set at p < 0.05.

### RNA-seq sample preparation

RNA samples extracted using RNeasy mini-Kit and RNase-free DNase (QIAGEN 74106). QC, library preparation, sequencing, and mapping were performed by Novogene using an Illumina Novaseq 6000 with 150bp paired-end reads.

### Proteomics LC-MS/MS

Proteomic analysis was conducted by liquid chromatography–tandem mass spectrometry (LC–MS/MS) at the Proteomics Service of the Servicio Central de Soporte a la Investigación Experimental (University of Valencia). Protein samples were collected in RIPA buffer under the same conditions used for Western blot analysis, and 10 µg of total protein was used per sample.

Sample preparation followed the SP3 protocol for detergent removal and sample clean-up prior to enzymatic digestion with trypsin, as described by Moggridge et al. (2018) and Müller et al. (2020), with minor modifications^29,30^. Following digestion, 2 µL of each sample were diluted to a final volume of 20 µL with 0.1% formic acid and loaded onto Evotip Pure tips (EvoSep) according to the manufacturer’s instructions. Peptide identification was carried out using MSFragger (via FragPipe) against the SwissProt human protein database. Protein quantification and differential expression analysis were performed using the default DIA-NN workflow.

### Bioinformatics analysis

RNA-seq data were quantified using the SeqMonk RNA-seq quantitation pipeline (http://www.bioinformatics.babraham.ac.uk) and normalized by total read count, expressed as reads per million mapped reads. Differential expression analysis was carried out using DESeq2, applying a cutoff of log₂ fold change ≥ 1 and adjusted p-value < 0.05, with multiple testing correction via the Benjamini–Hochberg method. For enhanced stringency, DESeq2 results were integrated with intensity difference filtering in SeqMonk.

Heatmaps and principal component analysis (PCA) plots were produced in SeqMonk. Gene ontology (GO) analysis was performed on significantly upregulated and downregulated genes, with enriched GO terms identified using DAVID^31^, applying a Benjamini-adjusted p-value threshold of < 0.05.

Raw label-free quantitative proteomics data were analyzed in R (version 2024.09.1) using the *Differential Enrichment analysis of Proteomics data* (DEP) package (version 1.26.0). Protein intensity values were imported from a pre-processed matrix containing LFQ (label-free quantification) intensities, previously generated using the DIA-NN protein quantification software.

Proteins with missing values in any sample were excluded by applying a strict missing-value threshold of zero, retaining only those quantified across all samples. The resulting dataset was normalized using variance stabilizing normalization (VSN) to minimize technical variation between samples. Differential enrichment analysis was then performed, with the control group explicitly defined. Proteins were considered significantly regulated if they exhibited an absolute fold change > 1.5 and an adjusted p-value < 0.05. Significant proteins were extracted based on these criteria.

BAP1-OE gene signature interrogation was performed as described in^32^, including unsupervised clustering and principal component analysis (PCA) to evaluate the ability of the gene list to separate the groups within each dataset. Hierarchical clustering heatmaps were generated using the gplots R package with Euclidean distance. Individual PCA plots were generated with the factoextra R package.

Single-sample Gene Set Enrichment Analysis (ssGSEA) was performed using the GSVA R package to compute enrichment scores (ES) for the BAP1-OE gene signature in individual samples. The signature was evaluated across placental transcriptomic datasets of early-onset preeclampsia (EO-PE; GSE114691 and GSE190971) and additional pregnancy complications associated with placental dysfunction, including intrauterine growth restriction (IUGR; GSE220877), gestational diabetes mellitus (GDM; GSE203346), and COVID-19 infection (GSE171995). Differences in ssGSEA ES between case and control groups were assessed using an unpaired two-tailed Student’s t-test. The diagnostic performance of the BAP1-OE signature in discriminating EO-PE from controls was evaluated using the pROC R package. ROC curves were generated from ssGSEA ES values, and the area under the curve (AUC), sensitivity, specificity, and optimal threshold (Youden index) were calculated.

For data exploration and visualization, PCA, volcano plots, hierarchical clustering heatmaps, and sample correlation plots were generated using a combination of DEP-native and custom R functions. Functional enrichment analysis of significantly regulated proteins was performed using Gene Ontology (GO) terms, and resulting p-values were transformed to –log₁₀ for graphical representation in bar plots. Protein–protein interaction networks were constructed using the STRING database (version 11).

### Quantification and statistical analysis

All statistical procedures, including sample sizes (*n*), number of independent experiments, and specific tests applied, are detailed in the Methods sections and in each figure legend. In brief, human placental sample comparisons (Fig. 1) were performed using Student’s two-tailed t-tests or one-way ANOVA, depending on the number of groups. Data from human samples are presented as median ± interquartile range, and sample sizes were limited by the availability of rigorously phenotyped placental material (Supplementary File 7). For cell-line experiments, each dataset represents ≥3 independent experiments. Unless otherwise specified, quantitative data from cell-line–based assays are presented as mean ± SEM. Proliferation analyses (Fig. 2D; Supplementary Fig. 2E) were performed using two-way ANOVA with Holm–Sidak post hoc correction. Adhesion assays (Fig. 2E; Supplementary Fig. 2F), western blot quantification (Fig. 1C, 2B–C, 5D; Supplementary Fig. 2C), and organoid area measurements (Supplementary Fig. 5B, 5F) were analysed using Student’s two-tailed t-tests, where n refers to the number of independent experiments or individual organoids, as specified in the figure legends. RT-qPCR and western blot analyses of PR-DUB complex dynamics during trophoblast differentiation were performed using one-way ANOVA with Dunnett’s multiple-comparisons test (Fig. 1A–B). RT-qPCR datasets quantifying lineage marker expression during EVT and STB differentiation (Fig. 3C, 4C; Supplementary Figs. 3A, 4A–B), HLA-G and SDC1 western blot quantification (Fig. 3B, 4B), and zymography (Supplementary Fig. 3B) were analysed using two-way ANOVA with Tukey’s multiple-comparisons test, with n representing the number of independent experiments.

**Figure 1.**
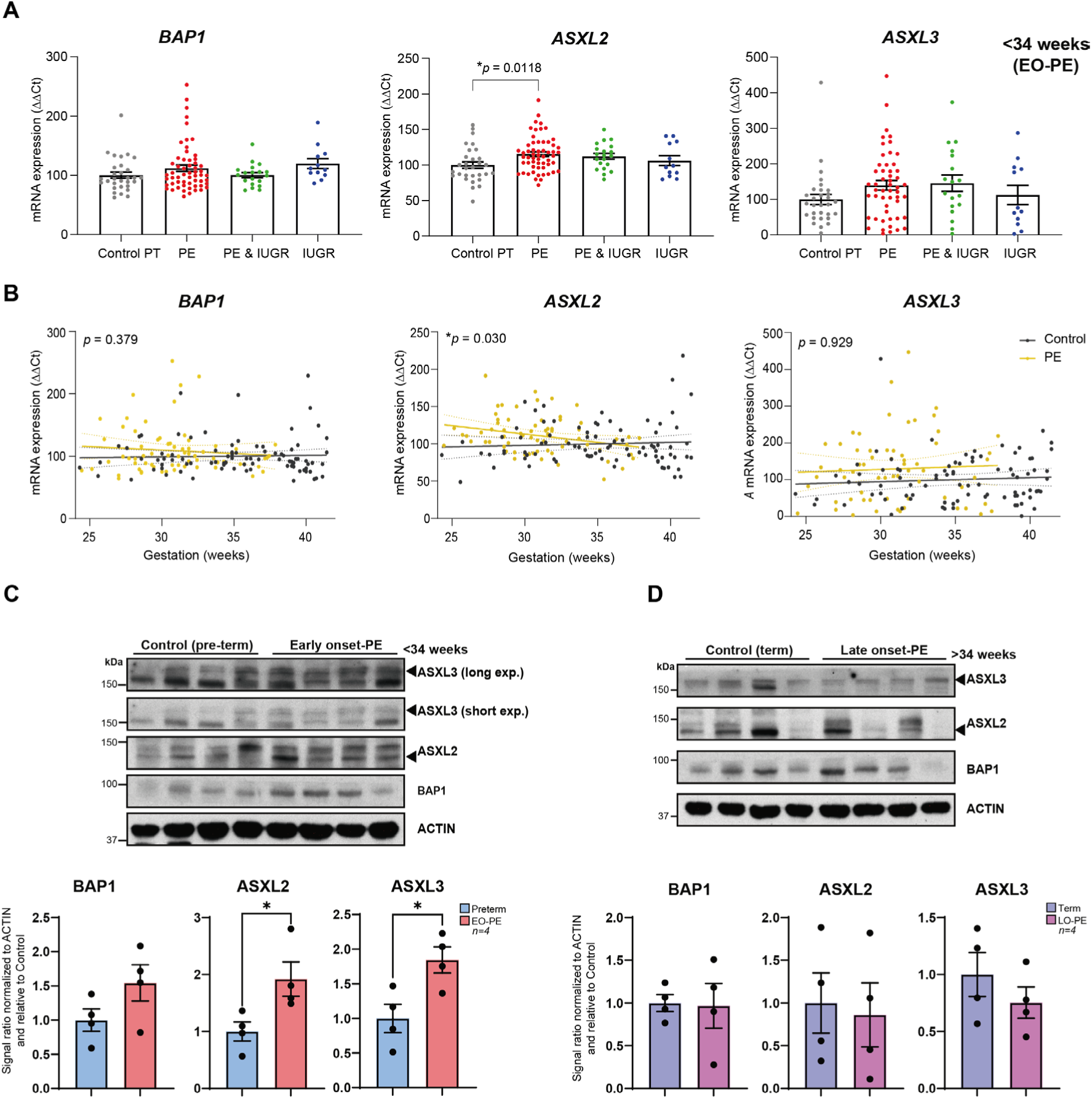
BAP1 PR-DUB complex levels in placental tissue from pregnancies with complications. **(A)** Relative mRNA expression of *BAP1*, *ASXL2*, and *ASXL3* in central villous placental biopsies collected before 34 weeks of gestation from gestational age–matched control pregnancies (Control PT; preterm, *n*=30), preeclampsia (PE, *n*=56), PE with intrauterine growth restriction (PE&IUGR, *n*=19), or IUGR alone (*n*=12). Data are presented as median ± interquartile range. Statistical analysis was performed using one-way ANOVA with Kruskal–Wallis multiple comparisons test. **(B)** mRNA levels of *BAP1*, *ASXL2*, and *ASXL3* were increased in early-onset PE (EO-PE), with *ASXL2* showing a significant association with gestational age at delivery (*p*=0.030). Associations were assessed using simple linear regression. *p* < 0.05. **(C)** Western blot analysis of BAP1, ASXL2, and ASXL3 protein levels in Control PT and EO-PE placentas (<34 weeks) from an independent cohort confirmed increased protein levels of the BAP1 PR-DUB complex in EO-PE. Graphs show quantification from four independent biological replicates. Data are presented as mean ± SEM; *p*<0.05 (Student’s *t*-test). **(D)** Western blot analysis of BAP1, ASXL2, and ASXL3 protein levels in late-onset PE (LO-PE) revealed no significant differences compared to Control PT. Graphs show quantification from four independent biological replicates. Data are presented as mean ± SEM (Student’s *t*-test).

**Figure 2:**
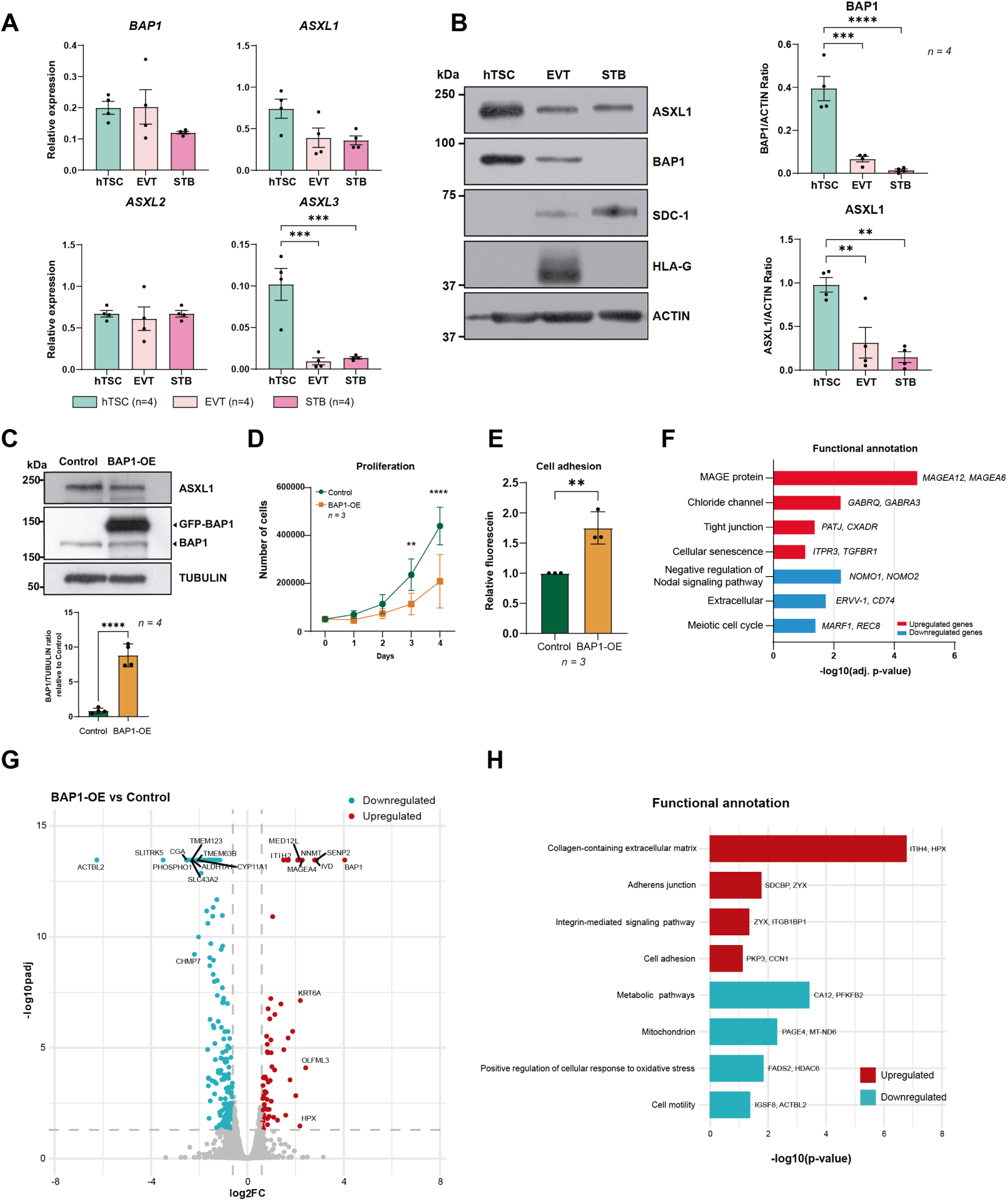
BAP1 downregulation is essential for early placental development. **(A)** mRNA expression levels of PR-DUB components (*BAP1, ASXL1, ASXL2,* and *ASXL3*) in human trophoblast stem cells (hTSCs), extravillous trophoblasts (EVT), and syncytiotrophoblasts (STBs), quantified by RT-qPCR. Data represent mean ± SEM of *n=*4 independent experiments; ***p<0.001; one-way ANOVA with Dunnett’s multiple comparisons test. (**B**) Western blot analysis of PR-DUB subunits (ASXL1, and BAP1) and lineage markers (SDC1 as a STB marker, HLA-G as an EVT marker) with ACTIN as a loading control, across differentiation states. Quantification of BAP1/ACTIN and ASXL1/ACTIN protein ratios are plotted on the right, confirming ASXL1 and BAP1 downregulation during differentiation. Data are mean ± SEM of *n=*4 independent experiments; **p<0.005, ***p<0.001, ****p<0.0001; one-way ANOVA with Dunnett’s multiple comparisons test. (**C**) Western blot of ASXL1, endogenous BAP1, GFP-tagged BAP1 (GFP-BAP1), and TUBULIN (loading control) in BAP1-OE and Control hTSCs. Quantification of BAP1/TUBULIN ratio is plotted on the bottom (mean ± SEM, *n*=4 independent experiments; ****p<0.0001; Student’s two-tailed t-test). (**D**) Proliferation rates of BAP1-OE and control hTSCs over 4 days, showing significant reduced cell numbers at days 3 and 4 (mean ± SEM, *n=*3 independent experiments; two-way ANOVA with Holm-Sidak’s post hoc test) (**E**) Adhesion assay quantifying fluorescein-labeled BAP1-OE hTSCs relative to controls (mean ± SEM, *n=*3 independent experiments; **p<0.005; Student’s two-tailed t-test). (**F**) Functional enrichment analysis of DEGs, highlighting pathways associated with upregulated (red) and downregulated (blue) genes. **(G**) Volcano plot of differentially expressed proteins (DEPs) in BAP1-OE hTSCs compared to control hTSCs (red: upregulated proteins; blue: downregulated proteins; adjusted *p*< 0.05, FC > 1.5). (**H**) Functional enrichment analysis of DEPs, highlighting pathways associated with upregulated (red) and downregulated (blue) proteins. OE, overexpression.

**Figure 3:**
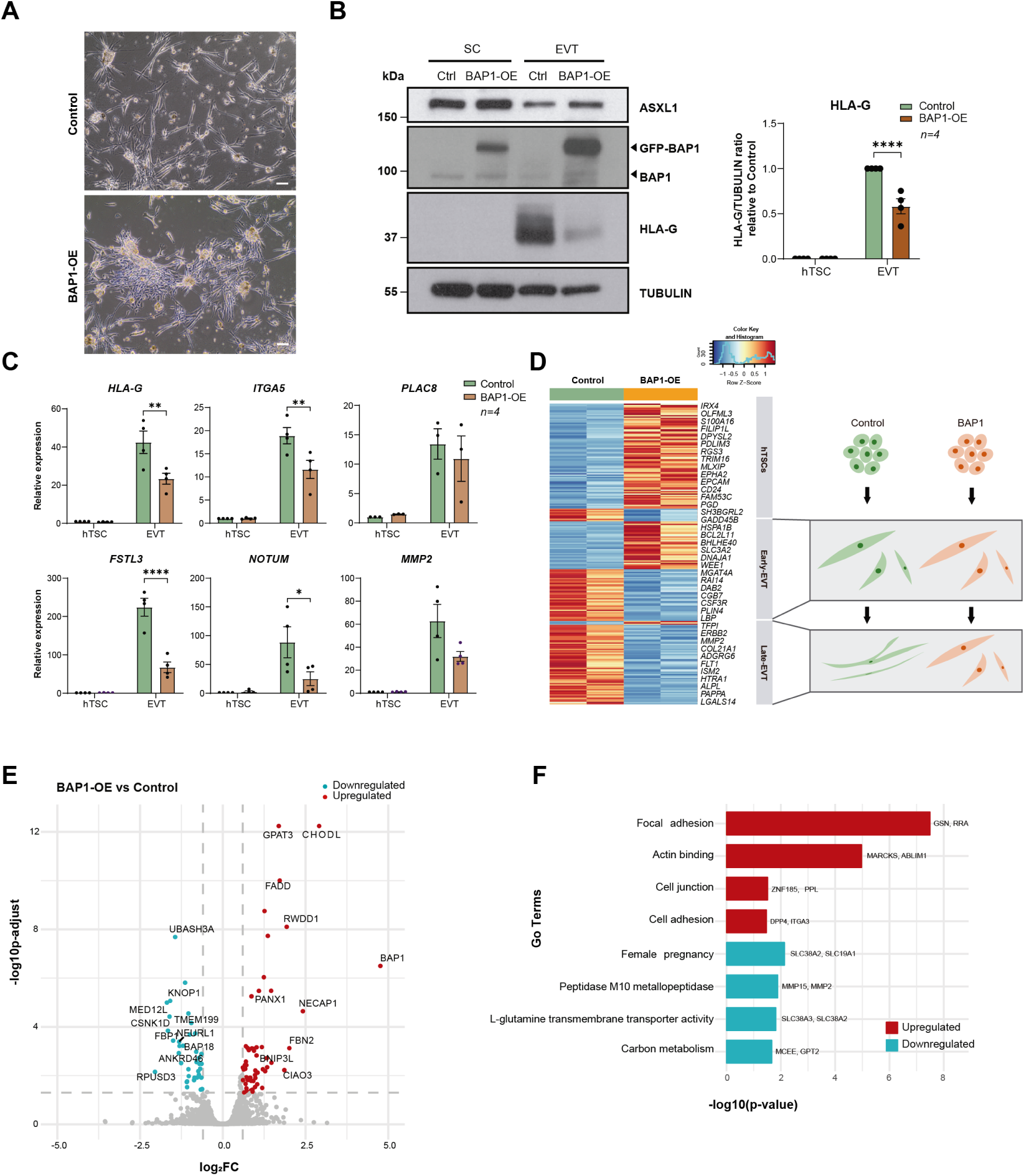
BAP1 overexpression impairs extravillous trophoblast formation and function. (**A**) Phase contrast images of EVTs showing aggregated colony morphology with reduced elongation in BAP1-OE cells versus control EVTs. Scale bars, 150 µm. (**B**) Western blot analysis of ASXL1, BAP1, GFP-BAP1, HLA-G and TUBULIN (loading control) in BAP1-OE hTSC and EVTs compared to control cells. Quantification of HLA-G/TUBULIN ratio relative to control cells growing in stem cell conditions is plotted on the right (mean ± SEM, *n=*4 independent experiments; ****p<0.0001; two-way ANOVA with Tukey’s multiple comparisons test. (**C**) RT-qPCR analysis of EVT-specific (*HLA-G, ITGA5, PLAC8, FSTL3, NOTUM, MMP2*) markers in BAP1-OE hTSC and EVTs compared to control cell (mean ± SEM, *n=*4 independent experiments; *p<0.05, **p<0.01, ****p<0.0001; two-way ANOVA with Tukey’s multiple comparisons test. (**D**) On the left, heatmap of stage-specific marker expression (row-scaled Z-scores) in BAP1-OE EVTs versus control EVTs, showing retention of early-EVT markers (*DAB2*, *LBP)* and failure to activate late-EVT (*MMP2*, *FLT1*) transcriptional programs. Gene clusters defined by Kim et al., 2024. On the right, schematic model of differentiation blockade, demonstrating how BAP1 overexpression prevents progression from early to late EVT states. (**E**) Volcano plot of differentially expressed proteins (DEPs) in BAP1-overexpressing EVTs compared control EVTS (red: upregulated proteins; blue: downregulated proteins; adjusted *p*< 0.05, FC > 1.5). (**F**) Gene ontology enrichment analysis of DEPs, highlighting pathways associated with upregulated (red) and downregulated (blue) proteins. EVT, extravillous trophoblast; hTSC, human trophoblast stem cell; OE, overexpression.

**Figure 4:**
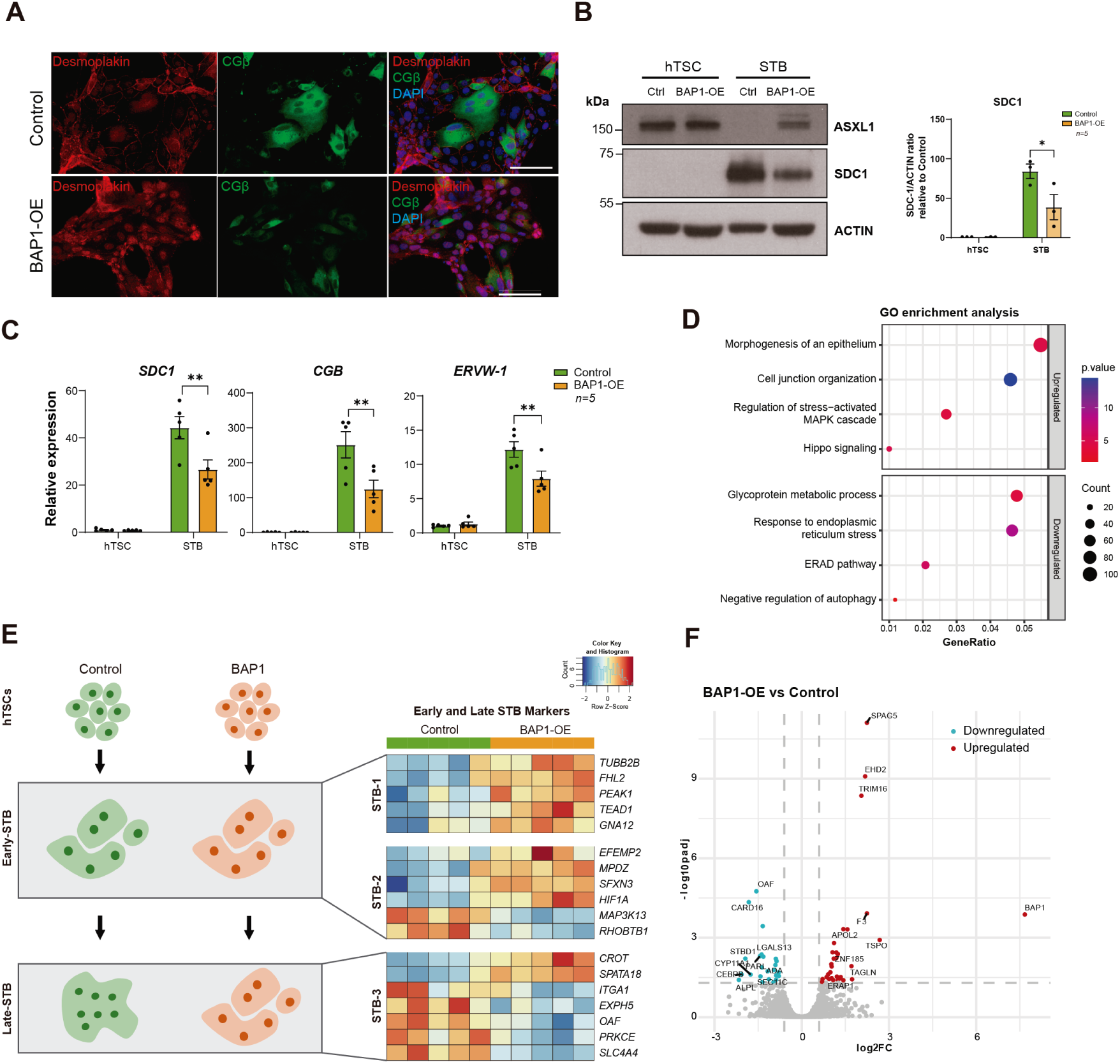
BAP1 overexpression impairs syncytiotrophoblast formation and function. (**A**) Immunofluorescence of Desmoplakin and CGβ in BAP1-OE and Control STBs, with DAPI nuclear counterstain. Scale bars, 150 µm. (**B**) Western blot of ASXL1, SDC1 and ACTIN (loading control) expression in BAP1-OE hTSC and STBs compared to control cells. Quantification of SDC1/ACTIN ratio relative to control hTSCs is plotted on the right (mean ± SEM, *n*=4 independent experiments; \**p*<0.05, two-way ANOVA with Tukey’s multiple comparisons test). (**C**) RT-qPCR analysis of STB-specific (*SDC1*, *CGB*, *ERVW-1*) markers in BAP1-OE hTSC and STBs compared to control cells (mean ± SEM of *n*=4 independent experiments; \*\**p*<0.005, two-way ANOVA with Tukey’s multiple comparisons test). (**D**) GO analysis of differentially expressed genes in BAP1-OE versus control STBs, highlighting upregulated and downregulated pathways. (**E**) On the left, a schematic of impaired cell fusion in BAP1-overexpressing cells, leading to defective STB formation. On the right, a heatmap of FPKM values for STB lineage markers (STB-1, STB-2, and STB-3), representing progressive differentiation states, from early to late STB states, in BAP1-OE and Control STBs. Data are z-score normalized across rows. (**F**) Volcano plot of differentially expressed proteins (DEPs) in BAP1-overexpressing STBs versus Controls (red: upregulated proteins; blue: downregulated proteins; adjusted *p*< 0.05, FC > 1.5). hTSC, human trophoblast stem cell; OE, overexpression; STB, syncytiotrophoblast.

RNA-seq differential expression analysis was performed using DESeq2, with significance defined as adjusted p < 0.05 (Benjamini–Hochberg correction) and |log₂FC| ≥ 1. PCA, volcano plots, hierarchical clustering, and GO analyses were generated using SeqMonk or R, as described (Fig. 2F, 3D, 4D–E, 5E; Supplementary Fig. 2J, 3C–E, 4C–H, 5G; Supplementary Files 1–2). Proteomics data were re-processed using DIA-NN for peptide quantification and the DEP R package for differential protein analysis. To minimize false positives, a stringent threshold of adjusted p < 0.05 and FC > 1.5 was applied across all proteomic datasets (Fig. 2G–H, 3E–F, 4F; Supplementary Fig. 4I, 5H–I; Supplementary Files 3–4). To define the BAP1-OE gene signature, only transcripts and proteins showing concordant differential regulation in both RNA-seq and proteomics were retained (Fig. 5F; Supplementary File 5). Enrichment of this signature in placental datasets was quantified using ssGSEA. ROC analyses were performed using the pROC R package to evaluate diagnostic performance in EO-PE versus controls (Fig. 5G; Supplementary Fig. 5J–K), with AUC, sensitivity, specificity, and optimal thresholds computed using the Youden index. All statistical analyses were performed using GraphPad Prism v9.4.1 and R (v2024.09.1). Statistical significance was defined as p < 0.05.

**Figure 5:**
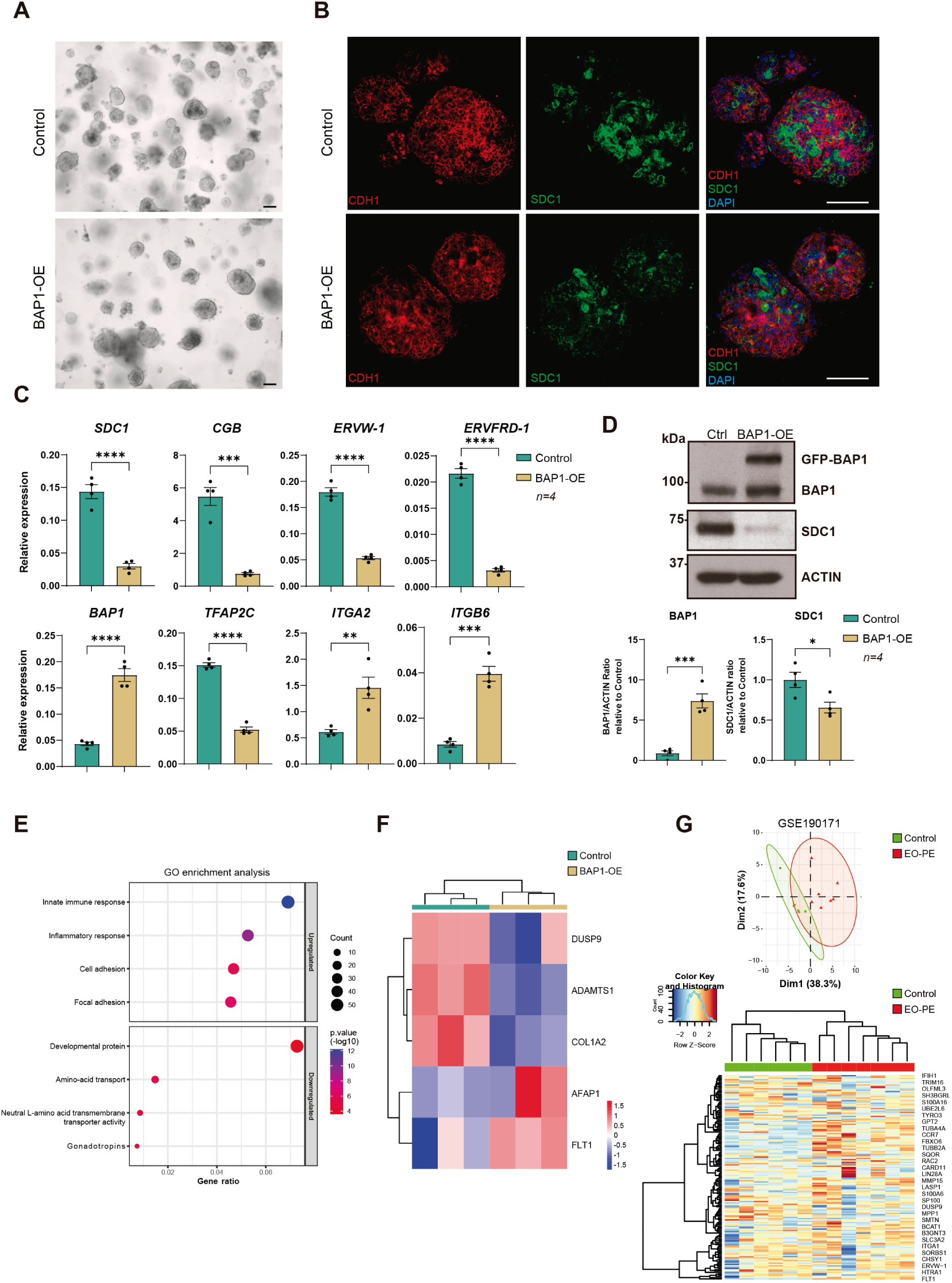
BAP1 overexpression disrupts trophoblast organoid formation and recapitulates early-onset preeclampsia features. (**A**) Brightfield images of BAP1-OE and control organoids, demonstrating comparable morphology and size. Scale bar, 150 um. (**B**) Immunofluorescence (IF) staining of BAP1-OE and control organoids for CDH1 and SDC1. Nuclear counterstaining was with DAPI. Scale bars, 150 µm. (**C**) RT-qPCR analysis of STB (*SDC1*, *CGB*, *ERVW*-*1*, and *ERVFRD-1*), *BAP1*, stem cell (*TFAP2C)* and villous cytotrophoblast cell column (*ITGA2*, *ITGB6*) markers in BAP1-OE and control organoids (mean ± SEM, *n=*4 independent experiments; \*\**p*<0.005, \*\*\**p*<0.001, \*\*\*\**p*<0.0001; Student two-tailed t-test). (**D**) Western blot analysis of BAP1, GFP-BAP1, SDC1, and ACTIN (loading control) in BAP1-OE and control organoids. Quantification of BAP1/ACTIN and SDC1/ACTIN ratio relative to control organoids is plotted on the bottom (mean ± SEM, *n=*3 independent experiments; *p<0.05, \*\*\**p*<0.001; Student’s two-tailed t-test). (**E**) GO enrichment analysis of differentially expressed genes (DEGs) in BAP1-OE *vs* control organoids. (**F**) Heatmap showing representative proteins from the BAP1-OE signature that are also dysregulated in PE, with downregulation of DUSP9, ADAMTS1, and COL1A2, and upregulation of AFAP1 and FLT1 in BAP1-OE organoids. Colour scale indicates scaled expression values (red, upregulation; blue, downregulation). (**G**) (Upper panel) Principal component analysis (PCA) plots showing the distribution of early-onset preeclampsia (EO-PE) samples (red) and controls (green) from the GSE190171 EO-PE-affected dataset, based on the BAP1-OE molecular signature. The X and Y axes represent principal components 1 and 2, which explain the indicated percentage of total variance. (Lower panel) Unsupervised hierarchical clustering of EO-PE (red) and control (green) samples from each dataset, based on the BAP1 gene signature. OE, overexpression; STB, syncytiotrophoblast.

## Data availability

The RNA-seq and proteomics datasets generated in this study have been deposited in publicly accessible repositories. RNA-seq data are available in the Gene Expression Omnibus (GEO), and proteomics data are available in the PRIDE database. Accession numbers for both datasets will be provided once the repositories complete the final assignment process (TBD). Raw data for all western blot experiments are included in Supplementary Material 1. All additional data supporting the findings of this study are available within the article, the Supplementary information files, or from the corresponding authors upon reasonable request.

## RESULTS

### BAP1 and PR-DUB components are upregulated in early-onset preeclampsia placentas

To explore whether BAP1 dysregulation is associated with preeclampsia, we analyzed placental tissues from two independent cohorts of pregnancies complicated by early-onset PE (EO-PE) and intrauterine growth restriction (IUGR). RT-qPCR analysis revealed an overall trend toward upregulation of *BAP1*, *ASXL2*, and *ASXL3* mRNA in EO-PE samples, with the increase in *ASXL2* reaching statistical significance, whereas *ASXL1* levels remained unaltered (Fig. 1A; Supplementary Fig. 1A). This expression profile was predominantly observed before 34 weeks of gestation (Fig.1B; Supplementary Fig 1B). Western blot analysis of an independent cohort corroborated these findings at the protein level: placentas from EO-PE cases exhibited elevated BAP1, ASXL2, and ASXL3 expression compared to gestationally matched control (control pre-term), whereas late-onset PE (LO-PE) samples showed no significant changes compared to gestationally matched control samples at term (Fig. 1C and 1D, quantifications below). These data indicate that BAP1-PR-DUB complex dysregulation is specifically associated with EO-PE.

### BAP1 regulates epithelial identity in human trophoblast stem cells

Previously, we have demonstrated that BAP1 is highly expressed in proliferative CTB within first-trimester human placentas but is markedly reduced in differentiated EVT and even further diminished in STB^24^. Given its dynamic expression during *in vivo* differentiation, we here examined the role of BAP1 PR-DUB complex in undifferentiated hTSCs and across hTSCs *in vitro* differentiation. RT-qPCR analysis showed that *ASXL1* was highly expressed in hTSCs under stem cell conditions and became markedly downregulated during trophoblast differentiation, both in EVT and STB. *ASXL3* mRNA levels also decreased upon differentiation, whereas BAP1 expression remained relatively stable, and *ASXL2* displayed only minor changes across differentiation. The expression patterns observed in EVT were consistent with our previous findings^24^, and here we extend these analyses to STB differentiation (Fig. 2A). At the protein level, ASXL1 was highly expressed under self-renewing conditions, while both BAP1 and ASXL1 levels declined upon EVT or STB differentiation (Fig. 2B). Knockdown of BAP1 via CRISPR/Cas9-mediated deletion of exon 4 in bulk BTS11 hTSCs led to increased proliferation and reduced cell adhesion, without compromising the stem cell marker profile (Supplementary Fig. 2A-G). Conversely, BAP1 overexpression (BAP1-OE) resulted in decreased proliferation and enhanced adhesion, with no gross morphological alterations (Fig. 2C-E; Supplementary Fig. 2H,I). To explore the molecular consequences of BAP1 overexpression, we performed RNA-seq analysis on both control and BAP1-OE hTSCs. This analysis identified 54 differentially expressed genes (DEGs) regulated by BAP1 overexpression (28 genes upregulated and 26 genes downregulated, *p-value*<0.05; log_2_FC>1) (Supplementary Fig. 2J; Supplementary file 1). In line with our previous observations of increased epithelial markers in BAP1-OE hTSC Gene Ontology (GO) analysis of BAP1-OE hTSCs revealed enrichment of gene programs related to cell adhesion, tight junction assembly, and epithelial integrity, alongside downregulation of cell cycle and metabolic regulators (Fig. 2F; Supplementary file 2).

Proteomic profiling (Differentially Expressed Proteins; DEP, FC>1.5, p-adjusted < 0.05) further confirmed that BAP1-OE cells exhibit increased abundance of proteins involved in the regulation of the collagen-containing extracellular matrix, adherens junction, integrin-mediated signaling pathways, and cell adhesion—hallmarks of reinforced epithelial traits—while downregulated proteins were enriched for functions related to metabolic pathways, mitochondrion, positive regulation of cellular response to oxidative stress and cell motility (Fig. 2G,H; Supplementary files 3 and 4). These findings indicate that BAP1 promotes an epithelial-like, adherent trophoblast state that is antagonistic to differentiation.

### Forced BAP1 expression delays EVT differentiation and invasion capacity

To assess the impact of BAP1 on EVT generation, we differentiated control and BAP1-OE, and BAP1-KD hTSCs for 8 days. Control and BAP1-KD cells successfully differentiated into EVTs, acquiring spindle-like mesenchymal morphology and upregulated canonical EVT markers, including *HLA-G*, *MMP2, ITGA5*, and *PLAC8*, with no significant differences between BAP1-KD and controls (Supplementary Fig. 3A). In contrast, BAP1-OE cells formed compact aggregates with attenuated HLA-G expression (Fig. 3A,B). RT-qPCR analyses demonstrated impaired induction of EVT markers, including *HLA-G*, *ITGA5*, *FSTL3*, *NOTUM*, and *MMP2* in BAP1-OE cells compared to control hTSCs (Fig. 3C). Zymography confirmed reduced MMP2 enzymatic activity (Supplementary Fig. 3B). Transcriptomic profiling (DESeq2, log_2_FC>1, *p-value* <0,05) revealed that BAP1-OE EVTs retained expression of progenitor and early-EVT markers while failing to activate genes associated with mature EVT identity and extracellular matrix remodeling^33^ (Fig. 3D; Supplementary Fig. 3C, D; Supplementary files 1 and 2). Differential proteomic analysis (FC > 1.5, p-adjusted < 0.05) identified 60 proteins significantly upregulated and 37 downregulated in BAP1-OE EVTs compared to control cells (Fig. 3E; Supplementary file 3). Gene Ontology analysis showed that upregulated proteins were primarily enriched in pathways related to focal adhesion, actin binding, cell junction organization, and cell adhesion. In contrast, the downregulated proteins were enriched in categories related to female pregnancy, L-glutamine transmembrane transporter activity and peptidase M10 metallopeptidase activity (Fig. 3F; Supplementary Fig. 3D; Supplementary file 4). Together, these results demonstrate that BAP1 overexpression perturbs EVT differentiation and impairs ECM remodeling.

### BAP1 overexpression impairs syncytiotrophoblast differentiation and function

To determine whether BAP1 also impairs STB formation, we induced syncytialization in BAP1-OE, BAP1-KD and control hTSCs under 2D conditions. Control and BAP1-KD cells displayed typical multinucleation and upregulated syncytial markers, including *SDC1*, *CGΒ*, *ERVW-1* and *ERVFRD-1* with no significant differences between BAP1-KD and controls, indicating that BAP1 downregulation does not impair STB differentiation (Supplementary Fig. 4A). In contrast, BAP1-OE cells retained epithelial morphology, exhibited diminished fusion, and showed impaired upregulation of STB genes (Fig. 4A-C; Supplementary Fig. 4B)). Notably, *ASXL1* and *ASXL3* transcript levels remained elevated in BAP1-OE cells, consistent with a stalled differentiation state (Supplementary Fig. 4B).

Transcriptomic analysis (DESeq2; log2FC>1, *p-value*<0.05) indicated downregulation of ER stress response, autophagy, pregnancy hormone regulation, and glycoprotein metabolism pathways, features typically enhanced during syncytialization, while epithelial junction, hippo signaling, focal adhesion, stem cell proliferation, and programs were upregulated (Fig. 4D; Supplementary Fig. 4C-H). Consistently, comparison of DEGs to known early- and late-STB markers^34^ showed that control STBs upregulated late-STB markers (*ITGA1, EXPH5, OAG, PRKCE,* and *SLC4A4*), whereas BAP1-OE cells remained in an early-STB state, characterized by expression of early-STB markers such as T*UBB2B, FHL2, PEAK1, TEAD1, GNA12, EFEMP2, MPDZ, SFXN3,* and *HIF1A* (Fig. 4E).

Comprehensive proteomic profiling revealed that BAP1-OE STBs displayed 31 significantly upregulated and 24 downregulated proteins compared to controls (FC > 1.5, adjusted *p*< 0.05; Fig. 4F; Supplementary file 3). GO analysis showed that upregulated proteins were enriched in pathways related to focal adhesion, actin binding, cell adhesion and cell-matrix organization, consistent with reinforcement of epithelial traits. In contrast, downregulated proteins were associated with mitochondrial inner membrane, protein folding in the endoplasmic reticulum, and metabolic pathway function (Supplementary Fig. 4I; Supplementary File 4). Collectively, these results demonstrate that BAP1 overexpression obstructs STB fate acquisition.

### BAP1-OE trophoblast organoids mirror preeclampsia features

Trophoblast organoids, which closely resemble first-trimester placental villi, provide a physiologically relevant model for studying human placental development^6,7^. These 3D structures consist of an outer layer of proliferative CTBs that spontaneously differentiate into STBs within the organoid core. To validate our findings, we cultured BAP1-OE, BAP1-KD, and control hTSCs as trophoblast organoids. Consistent with findings in 2D cultures, BAP1-KD organoids exhibited larger size than controls, indicative of enhanced proliferation (Supplementary Fig. 5A, B). BAP1-KD organoids displayed defects in STB differentiation, including reduced SDC1 staining, disorganized CDH1 localization in the CTB layer, and downregulation of canonical STB markers (*SDC1*, *CGB*) (Supplementary Fig. 5C, D), suggesting compromised epithelial integrity and altered STB differentiation timing. BAP1-OE organoids also exhibited markedly reduced central syncytial regions (Fig. 5A, B; Supplementary Fig. 5E, F). Immunofluorescence and RT-qPCR analyses revealed downregulation of STB markers, including *SDC1, CGB, ERVW1,* and *ERVFRD-1*, alongside increased expression of CTB-associated integrins *ITGA2* and *ITGB6* (Fig. 5B, C; Supplementary Fig. 5E, F). Western blot analysis further confirmed impaired STB differentiation, as BAP1-OE organoids failed to upregulate SDC1, a key mediator of syncytial maturation (Fig. 5D).

RNA-seq revealed a profound impact of BAP1 overexpression on the organoid transcriptome, identifying 1936 differentially expressed genes (DESeq2; log₂FC > 1, *p* < 0.05), with 972 upregulated and 964 downregulated transcripts (Supplementary File 1). Principal component analysis (PCA) showed clear segregation of samples based on intracellular BAP1 levels (Supplementary Fig. 5G; Supplementary File 1). Gene ontology (GO) enrichment analysis indicated that upregulated genes were enriched for innate immune and inflammatory responses, focal adhesion, and cell adhesion pathways. In contrast, downregulated genes were associated with developmental protein, amino acid transport, neutral L-amino acid transmembrane transporter activity, and gonadotropin expression, features typically linked to impaired STB maturation (Fig. 5E, Supplementary File 2).

Complementary proteomic profiling identified 102 significantly upregulated and 63 downregulated proteins in BAP1-OE organoids (FC > 1.5, adjusted p< 0.05; Supplementary File 3). Reactome Pathway and GO analysis using STRING revealed enrichment of interferon signaling, cytokine signaling, MHC class I peptide folding and loading, ER-phagosome, and endosomal and vacuolar trafficking (Supplementary Fig. 5H). Proteins related to extracellular matrix organization, extracellular region and space, and endoplasmic reticulum lumen were significantly downregulated, reinforcing a disruption in STB differentiation (Supplementary Fig. 5I).

Integrated transcriptomic and proteomic analyses demonstrate that BAP1 overexpression induces a pro-inflammatory, progenitor-like state that is incompatible with proper syncytiotrophoblast maturation, a phenotype that mirrors key pathological features of EO-PE, including heightened interferon signaling and dysregulated immune–trophoblast interactions^35^. To further investigate this association, we defined a BAP1-OE gene signature by integrating transcriptomic and proteomic data across STB, EVT, and trophoblast organoid models. Differentially expressed genes (DEGs; DESeq2, log₂FC > 1, adjusted p< 0.05) and differentially expressed proteins (DEPs; FC > 1.5, p< 0.05) between control and BAP1-OE conditions showing concordant directional changes were retained (Supplementary File 5), providing a robust representation of BAP1-driven trophoblast states. When applied to trophoblast organoids, the BAP1-OE signature revealed dysregulation of genes previously linked to PE, including downregulation of DUSP9, ADAMTS1, and COL1A2, and upregulation of AFAP1 and FLT1 (Fig. 5F)^36–40^. The BAP1-OE gene signature was next interrogated across two independent EO-PE placental datasets (GSE114691 and GSE190971), as well as datasets representing other pregnancy complications associated with placental dysfunction, including intrauterine growth restriction (IUGR; GSE220877), gestational diabetes mellitus (GDM; GSE203346), and COVID-19 (GSE171995). In EO-PE cohorts, the BAP1 signature achieved an average sensitivity of 48.5%, specificity of 97.5%, and a mean AUC of 0.71, indicating strong discriminatory power with minimal false-positive classification. In contrast, the signature showed limited or no discriminatory capacity in IUGR, GDM, or COVID-19 placentas, underscoring its specificity for EO-PE–related placental alterations (Supplementary File 5). Principal component analysis (PCA) and unsupervised hierarchical clustering further confirmed a clear segregation of EO-PE from control samples, whereas other pregnancy disorders displayed overlapping transcriptional profiles (Fig. 5G; Supplementary Fig. 5J, K). Together, these findings indicate that the BAP1-driven molecular program constitutes a distinct feature of EO-PE placentas and provide a mechanistic link between BAP1 dysregulation, impaired STB lineage commitment, and the pathogenesis of early-onset preeclampsia.

## Discussion

This study identifies BAP1 as a central regulator of human trophoblast differentiation, with direct implications for placental development and the pathogenesis of early-onset preeclampsia (EO-PE). By integrating transcriptomic, proteomic, and functional data across 2D and 3D models, we demonstrate that BAP1 must be downregulated to enable lineage commitment toward both extravillous trophoblast (EVT) and syncytiotrophoblast (STB) fates. In contrast, sustained BAP1 expression maintains a progenitor-like, epithelial state and impairs terminal differentiation, providing a mechanistic link to the trophoblast dysfunction characteristic of EO-PE placentas.

Previous studies in mice have shown that Bap1 deletion causes mid-gestation lethality due to placental defects, including impaired chorionic ectoderm differentiation and failure of labyrinth formation, which compromises maternal–fetal exchange^20,41^. These defects have been linked to abnormal syncytiotrophoblast development and an excess of trophoblast giant cells, indicating a role for BAP1 in trophoblast lineage specification. At the molecular level, downregulation of BAP1 in mouse trophoblast stem cells promotes trophoblast epithelial–mesenchymal transition (EMT) characterized by reduced epithelial features, acquisition of mesenchymal and invasive traits, and enhanced trophoblast invasiveness^24^. However, whether this function is conserved in humans and whether it contributes to pregnancy complications has remained unclear.

Here, we show that BAP1 is highly expressed in human trophoblast stem cells (hTSCs) and undergoes robust downregulation during differentiation toward both EVT and STB lineages. Gain- and loss-of-function experiments reveal that persistent BAP1 expression stabilizes epithelial identity, enhances cell adhesion, and represses transcriptional programs required for lineage progression. These findings are consistent with BAP1’s role in maintaining epithelial organization in other tissues, such as the liver^42^, and underscore the broader role of BAP1 in regulating epithelial plasticity.

The biological effects of BAP1 dysregulation are strongly context-dependent. In most cancer models, including uveal and cutaneous melanoma as well as renal cell carcinoma, BAP1 loss promotes stemness, activation of EMT programs, and aggressive tumor behavior^43–46^. However, a recent study in uveal melanoma reported the opposite effect, showing that BAP1 loss drives epithelial gene expression and promotes cell aggregation^47^. These apparently conflicting results highlight the complexity of BAP1 function, which likely depends on the cellular and epigenetic context. Our findings expand this framework to the trophoblast lineage, where sustained BAP1 expression similarly stabilizes epithelial features and impedes both invasive and syncytial differentiation. This emphasizes that timely BAP1 downregulation is essential for normal trophoblast plasticity and placental development.

By modulating epithelial traits, EMT and cell invasion, BAP1 provides a mechanistic bridge between placental development and oncogenic transformation, as comparable molecular networks govern trophoblast invasion and tumour metastasis^48^. Although a direct role for BAP1 in EMT regulation during development or tumorigenesis remains incompletely defined, reduced BAP1 expression in cancers with prominent EMT components, such as uveal melanoma, clear-cell renal cell carcinoma, gastric adenocarcinoma, colorectal cancer, and non-small-cell lung cancer correlates with increased invasiveness and poorer prognosis^49–52^. These observations support a model in which low BAP1 levels facilitate EMT-driven malignancy. Nevertheless, the consequences of BAP1 loss vary across tissues. In renal tumor cells, for example, its inactivation promotes a mesenchymal-to-epithelial transition^53^, underscoring the cell–type–specific nature of its regulatory functions. Further studies will be needed to delineate how BAP1 interacts with lineage-specific transcriptional and chromatin programs to fine-tune EMT regulation in different biological contexts.

Considering that BAP1 downregulation appears critical for trophoblast plasticity, we investigated how altering BAP1 expression, either through overexpression or knockdown, perturbs the differentiation of trophoblast lineages *in vitro*. In monolayer cultures, BAP1 overexpression impaired EVT differentiation by preventing the induction of key markers such as *HLA-G*, *ITGA5*, and *MMP2*, and by reducing MMP2 enzymatic activity, indicating diminished invasive capacity, a hallmark of EO-PE pathophysiology ^10,53–55^. Under STB differentiation conditions, BAP1-overexpressing cells exhibited reduced *CGB* expression and formed fewer and smaller syncytial clusters. In contrast, BAP1 knockdown did not significantly impact EVT or STB differentiation in 2D monolayers, suggesting that basal BAP1 expression is sufficient to sustain normal lineage specification in these simplified systems. Bulk transcriptomic and proteomic analyses of BAP1-overexpressing cells confirmed a failure to activate gene networks involved in glycoprotein biosynthesis, autophagy, and ER stress responses—processes essential for STB maturation and hormone production, and frequently altered in EO-PE placentas^32,56,57^.

These functional defects were faithfully recapitulated in 3D trophoblast organoids, which model early villous architecture. Control organoids exhibited the expected organization, with a basal CTB layer surrounding a central STB core, whereas BAP1-overexpressing organoids showed reduced syncytialization, persistent expression of progenitor markers, and repression of endocrine differentiation pathways. Conversely, BAP1 knockdown produced an opposite yet complementary phenotype, characterized by excessive organoid growth and expansion of the CTB compartment, suggesting increased proliferative activity. Despite this hyperproliferation, BAP1-deficient organoids displayed defective STB formation, likely resulting from impaired spontaneous differentiation of the central cell population or disrupted signaling between CTB and emerging STB layers. This phenotype, where BAP1 depletion does not affect 2D differentiation but compromises STB generation specifically in 3D organoids, is consistent with previous observations^58^, emphasizing the importance of spatial organization and intercellular communication in regulating trophoblast maturation, and highlighting the more physiological relevance of 3D organoid systems compared to conventional 2D cultures^59–62^. Together, these findings indicate that both excessive and insufficient BAP1 activity disturb the balance between trophoblast proliferation and differentiation, highlighting the need for precise temporal and spatial control of BAP1 during villous development. Transcriptomic and proteomic analyses of BAP1-overexpressing organoids further revealed enrichment of inflammatory and immune-related pathways, including interferon signaling, MHC class I peptide-loading, and cytokine activity, suggesting that BAP1 overexpression not only blocks differentiation but also promotes immune dysregulation. This phenotype mirrors the inflammatory milieu observed in EO-PE placentas, marked by excessive type I interferon activity, altered innate immune system, and aberrant maternal–fetal immune crosstalk^35,63,643563^. These findings support a model in which BAP1 overexpression traps trophoblasts in an undifferentiated state, compromising both endocrine function and tissue remodeling capacity.

Mechanistically, BAP1 functions as the catalytic subunit of the PR-DUB complex, which removes monoubiquitin from histone H2A lysine 119 (H2AK119ub1), a modification associated with transcriptional repression^65^. PR-DUB function requires interaction with ASXL family proteins (ASXL1–3), which confer target specificity and modulate chromatin accessibility^66^. We show that expression of *BAP1*, *ASXL1* and ASXL3, decreases, consistent with the progressive loss of progenitor identity. In contrast, *BAP1*, *ASXL2*, and *ASXL3* are upregulated in EO-PE placentas from EO-PE cases before 34 weeks of gestation, correlating with disease severity. These findings suggest that dysregulation of PR-DUB activity may contribute to EO-PE by maintaining trophoblasts in a poorly invasive, poorly endocrine, progenitor-like state.

The upstream signals responsible for BAP1 and PR-DUB upregulation in early-onset preeclampsia (EO-PE) remain to be defined. Current evidence indicates that BAP1 expression and activity are dynamically modulated by cellular stress pathways, particularly hypoxia, whereas ER stress and interferon signaling are likely downstream consequences of excessive BAP1 activity rather than primary inducers. Under hypoxic conditions, BAP1 levels increase at both the mRNA and protein level, as demonstrated in nasopharyngeal carcinoma cells where hypoxia-induced BAP1 enhances erastin-induced ferroptosis through histone H2A deubiquitination and repression of SLC7A11^6667,68^. BAP1 also interacts with and stabilizes HIF-1α, promoting its nuclear accumulation and transcriptional activity, whereas BAP1 loss reduces HIF-1α levels and impairs adaptation to oxygen deprivation^69^. These findings suggest that hypoxia is a physiological inducer of BAP1 expression, potentially linking placental oxygen tension to chromatin-based regulation of trophoblast differentiation.

In contrast, ER stress and interferon signaling appear to be modulated by BAP1 itself. BAP1 binds to the promoters of unfolded protein response (UPR) genes such as ATF3 and CHOP, repressing their transcription during metabolic or ER stress ^70^. This regulatory mechanism normally serves to fine-tune the UPR, preventing excessive activation and apoptosis. However, our findings indicate that excessive BAP1 expression in trophoblasts excessively represses the UPR, thereby impairing the physiological ER stress response required for proper syncytialization^71^. In this context, BAP1 overexpression suppresses key UPR components and disrupts the adaptive stress signaling needed for the differentiation and fusion of villous cytotrophoblasts into syncytiotrophoblasts. This aberrant repression may contribute to defective syncytialization and placental dysfunction characteristic of EO-PE. Furthermore, BAP1 functions as an upstream activator of type I interferon pathways, inducing IFN-β expression and promoting STING and ISGF3 activation. Conversely, BAP1 loss weakens interferon signaling and reduces interferon-mediated tumor suppression^72^. These findings are consistent with our observation that BAP1-overexpressing trophoblast organoids display enhanced interferon signaling and a pro-inflammatory transcriptional phenotype that mirrors EO-PE. Together, these results suggest that aberrant BAP1 expression simultaneously suppresses adaptive ER stress responses and amplifies inflammatory interferon signaling, creating a dual imbalance that disrupts trophoblast homeostasis.

Collectively, this study identifies BAP1 as a key integrator of hypoxic, metabolic, and immune cues through chromatin-based regulation of gene expression. In the context of EO-PE, aberrant BAP1 activation may not only impair trophoblast differentiation but also exacerbate cellular stress and inflammatory responses, further compromising placental function. However, elevated BAP1 levels *in vivo* could initially arise as a secondary response to hypoxia, a hallmark of EO-PE placentas. Disentangling these causal relationships will require temporal analyses to determine how HIF-1α, UPR, and interferon pathways converge on BAP1 regulation in physiological versus pathological conditions.

In addition, our data implicate BAP1 in disrupting maternal–fetal immune tolerance. The suppression of HLA-G and concomitant activation of interferon-stimulated genes in BAP1-overexpressing trophoblasts mirror the immune alterations typical of EO-PE, where defective HLA-G expression and aberrant uterine NK cell function are hallmarks of disease^73,74^.

Determining whether BAP1 dysregulation represents a primary pathogenic event or a downstream consequence of placental stress will be essential to elucidate the sequence of events leading to placental failure. Ultimately, identifying the upstream triggers and downstream consequences of aberrant PR-DUB activity may yield novel biomarkers for early EO-PE detection and inform molecular strategies to restore trophoblast homeostasis. Together, our findings position BAP1 as a central gatekeeper of trophoblast differentiation and placental health, offering a mechanistic framework for understanding how disruptions in chromatin regulation contribute to the pathogenesis of preeclampsia.

## Limitations of the study

Although our data demonstrate that altered BAP1 levels disrupt trophoblast differentiation and recapitulate key molecular signatures of EO-PE, several limitations should be acknowledged. First, while we show that BAP1 and its cofactors are upregulated in EO-PE placentas, it remains unclear whether this reflects a primary pathogenic mechanism or a secondary adaptive response to placental stressors such as hypoxia, oxidative stress, or inflammation. Second, although we employ both BAP1 overexpression and knockdown models to interrogate the role of BAP1 in trophoblast biology, these perturbation systems may not fully capture the subtler and possibly heterogeneous dysregulation occurring *in vivo*. Future studies using patient-derived trophoblast organoids, primary cultures, or *in vivo* models will be essential to establish the causal hierarchy and physiological relevance of BAP1 dysregulation during placental development. Third, while our transcriptomic and proteomic datasets identify inflammatory and interferon-related pathways as major downstream consequences of BAP1 perturbation, we have not yet determined whether pharmacological or genetic inhibition of these pathways can restore normal trophoblast differentiation. Finally, although our study defines a global molecular landscape of BAP1-driven dysregulation, the specific mechanistic contributions of individual genes and proteins to EO-PE pathogenesis remain to be elucidated.

Addressing these limitations will be critical to delineate the upstream regulators and downstream effectors of BAP1 activity in the placenta and to evaluate its potential as a biomarker and therapeutic target for early-onset preeclampsia.

## Availability of Data and Materials

All data generated or analysed during this study are included in the manuscript and supporting files. Genome-wide sequencing data have been deposited in the GEO database under accession number TBD. The mass spectrometry proteomics data have been deposited in the ProteomeXchange Consortium via the PRIDE partner repository with the dataset identifiers TBD. Original western blot images have been deposited at Mendeley and are publicly available as of the date of publication. This paper does not report original code. Any additional information required to reanalyze the data reported in this paper is available from the corresponding author upon request.

## Acknowledgements

We thank Dr. Hiroaki Okae and Dr. Takahiro Arima for generously providing the hTSC cell lines used in this study. We are also deeply grateful to the donors whose contributions made this research possible. We acknowledge the support of the Advanced Light Microscopy Facility at the Centro de Biología Molecular “Severo Ochoa” (Madrid, Spain). This work was funded by the Ministerio de Ciencia e Innovación, Agencia Estatal de Investigación (MICIU/AEI/10.13039/501100011033), under grant numbers PID2020-114459RA-I00, PID2023-148535OB-I00, and CNS2022-135933, as well as by the Fundación Ramón Areces. A.F.-M. is supported by a PhD fellowship from the Asociación Española Contra el Cáncer (AECC) – Comunitat Valenciana. L.Y. was supported by the BITRECS fellowship programme (“Biomedicine International Training Research Programme for Excellent Clinician-Scientist”) at Hospital Clínic de Barcelona, funded by the “la Caixa” Foundation (grant code LCF/PR/SP23/52950012). F.C. received support from Hospital Clínic de Barcelona (Intensificació Interna) and the Fundació Jesús Serra of the Grup Catalana Occident.

## Author contributions

V.P.-G. conceptualized the study. P.D.-B., A.F.-M, S.N-S, M.M-L, J.G and C.M performed experiments and generated data for this study. LY, C.M, F.C and T.K undertook human placenta sample isolation. P.D.-B., and S.N-S performed bioinformatics analyses. V.P.-G drafted the manuscript, with input from all authors.

---

**Supplementary Figure 1:**
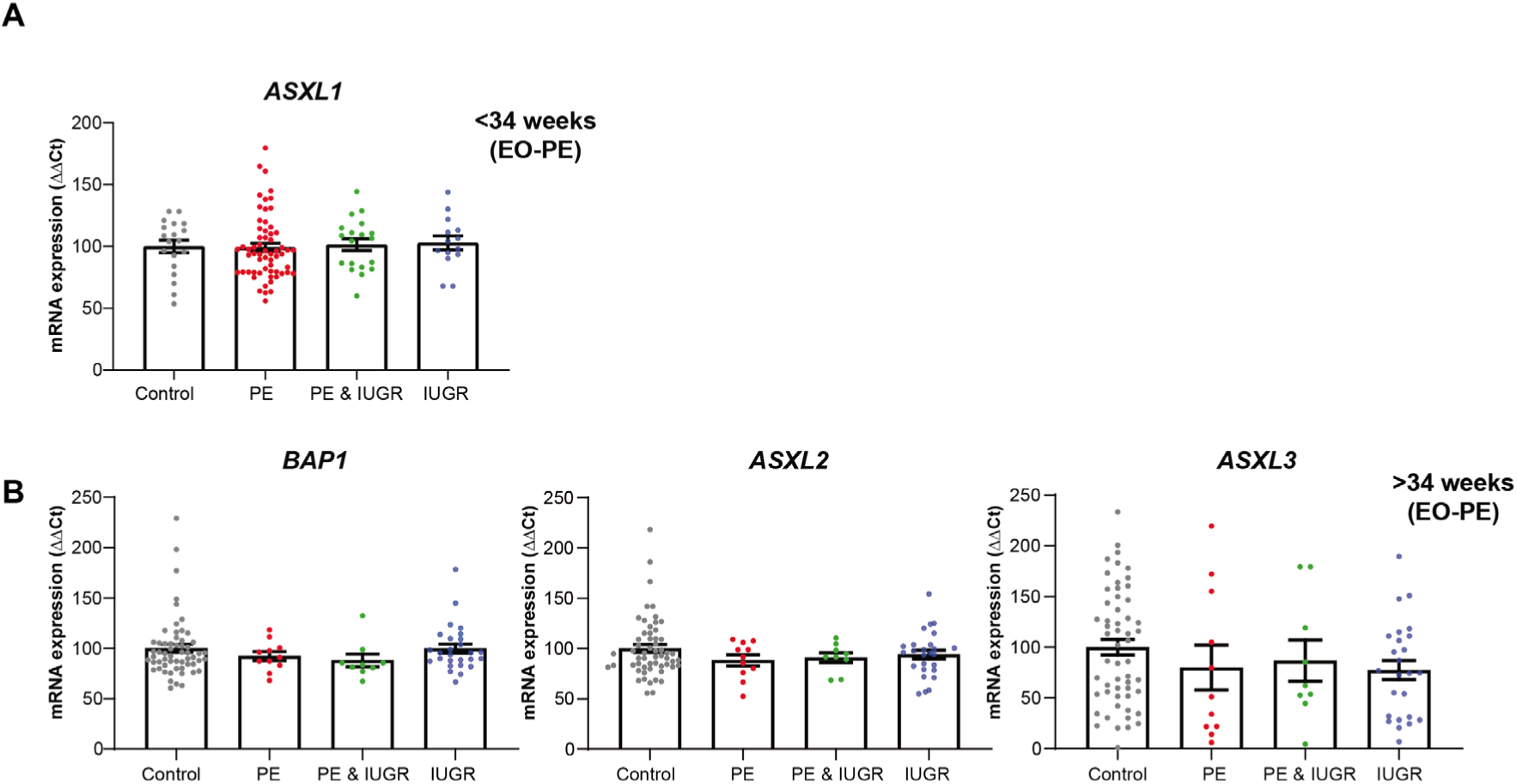
BAP1 PR-DUB complex levels in placental tissue from pregnancies with complications. **(A)** Relative mRNA expression of *ASXL1* in central villous placental biopsies collected before 34 weeks of gestation from gestational age–matched control pregnancies (Control PT; preterm, *n*=30), preeclampsia (PE, *n*=56), PE with intrauterine growth restriction (PE&IUGR, *n*=19), or IUGR alone (*n*=12). **(B)** Relative mRNA expression of BAP1, *ASXL2* and *ASXL3* in central villous placental biopsies collected after 34 weeks of gestation from gestational age–matched control pregnancies (Control; *n*=57), preeclampsia (PE, *n*=11), PE with intrauterine growth restriction (PE&IUGR, *n*=8), or IUGR alone (*n*=27).

**Supplementary Figure 2:**
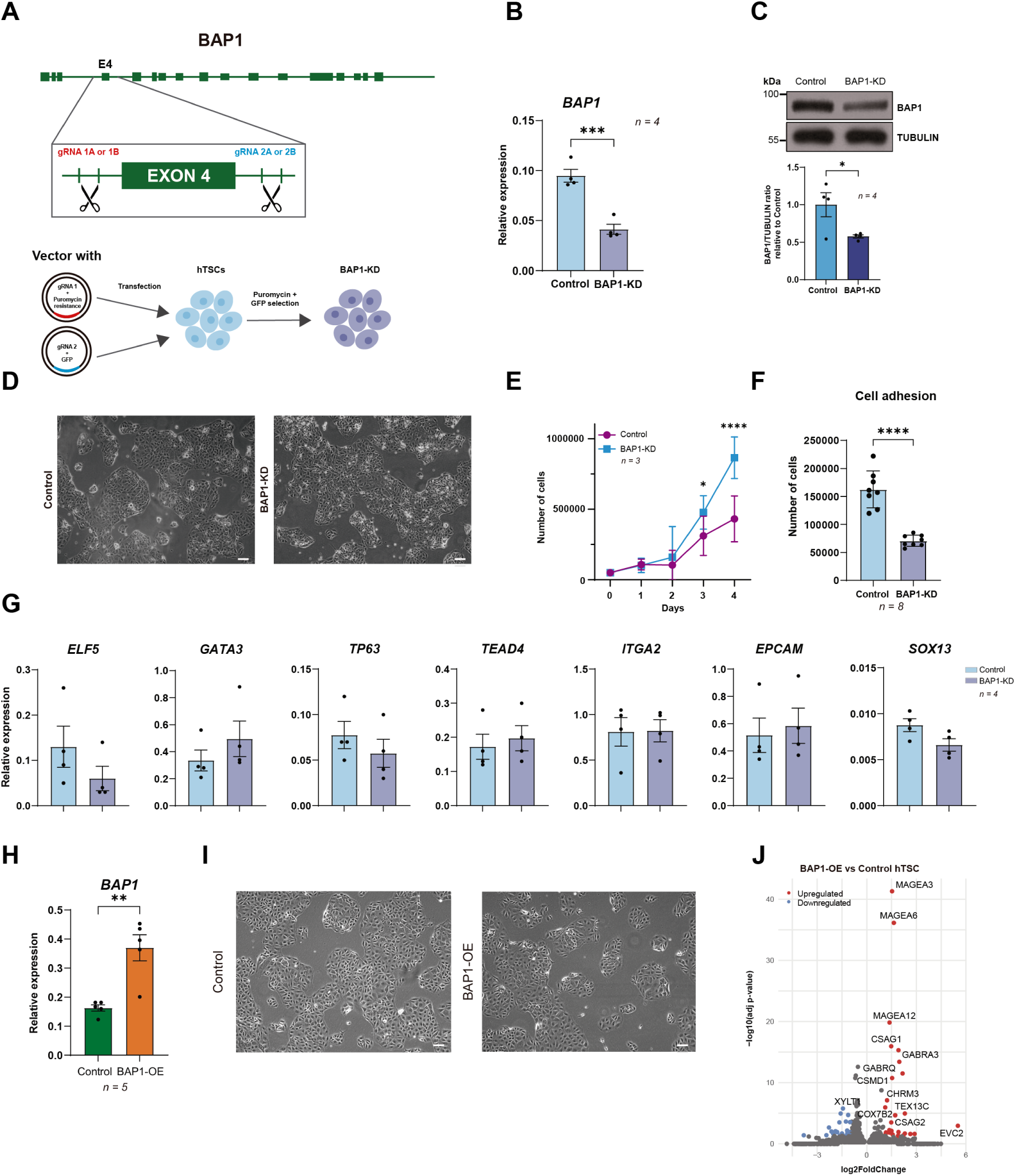
BAP1 knockdown and BAP1 overexpression alter trophoblast cell biology. (**A**) CRISPR-Cas9 targeting strategy for generating BAP1-knockdown (KD) hTSCs, showing guide RNA (gRNA) positioning in introns surrounding exon 4. (**B**) RT-qPCR analysis of *BAP1* (exon4) levels in BAP1-KD and control hTSCs (mean ± SEM, *n=*4 independent experiments; ***p<0.001; Student’s two-tailed t-test. (**C**) Western blot analysis of BAP1 and loading control (TUBULIN) in BAP1-KD versus Control hTSCs. Quantification of BAP1/TUBULIN ratio relative to controls is plotted on the bottom (mean ± SEM, *n=*4 independent experiments; *p<0.05; Student’s two-tailed t-test). (**D**) Phase-contrast images demonstrating conserved morphology in BAP1-KD *vs* Control hTCSs. Scale bars = 150µm. (**E**) Proliferation rates of BAP1-KD and control hTSCs over 4 days. BAP1 depletion significantly increased cell numbers at days 3 and 4 (mean ± SEM, *n=*3 independent experiments; *p<0.05, ****p<0.0001; two-way ANOVA followed by Holm-Sidak’s post hoc test). (**F**) Adhesion assay quantifying attached BAP1-KD and control hTSCs (mean ± SEM, *n*=8 independent experiments; ****p<0.0001, Student’s two-tailed t-test). (**G**) RT-qPCR analysis of and stem cell (*ELF5, GATA3, TP63, TEAD4, SOX13*) and epithelial (*EPCAM, ITGA2*) markers in BAP1-KD versus control hTSCs. Data represent mean ± SEM (*n=*4 independent experiments; Student’s two-tailed t-test). (**H**) RT-qPCR analysis of *BAP1* expression in BAP1-OE versus Control hTSCs (mean ± SEM, *n*=5 independent experiments; **p<0.01; Student’s two-tailed t-test). (**I**) Phase-contrast images confirming conserved morphology in BAP1-OE hTSCs. Scale bar = 150 µm. (**J**) Volcano plot of differentially expressed genes (DEGs) in BAP1-OE hTSCs versus controls (red: upregulated genes; blue: downregulated genes; adjusted *p*< 0.05, |log2FC| > 1). hTSC, human trophoblast stem cell; OE, overexpression.

**Supplementary Figure 3:**
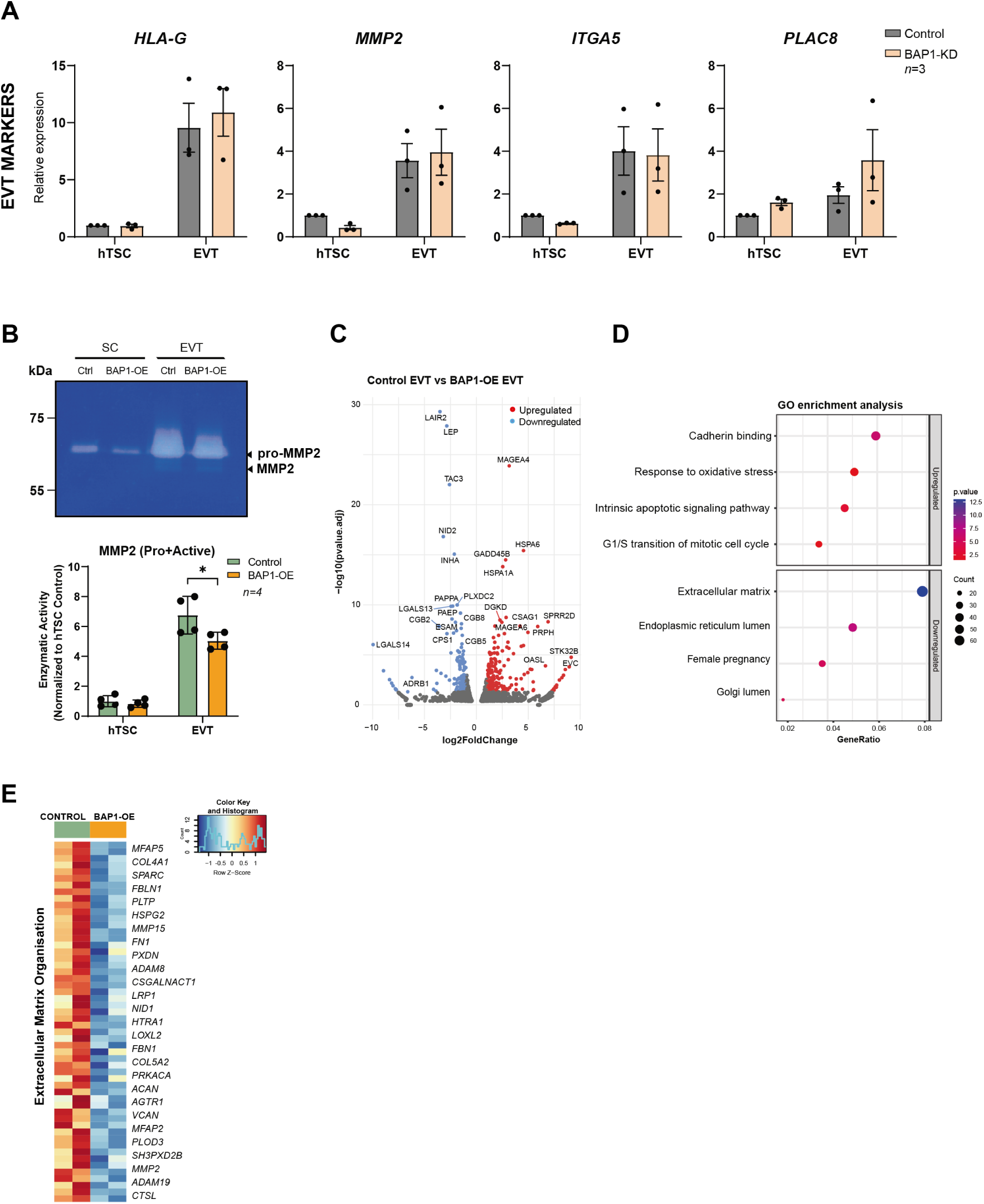
BAP1 overexpression impairs extravillous trophoblast differentiation. (**A**) RT-qPCR analysis of extravillous trophoblast (EVT) markers (*HLA-G*, *MMP2, ITGA5*, and *PLAC8*) in BAP1-KD human trophoblast stem cells (hTSC) and EVTs compared to controls. Data are mean ± SEM, *n=*3 independent experiments, two-way ANOVA with Tukey’s multiple comparisons test. (**B**) Gelatin zymography demonstrating reduced MMP2 proteolytic activity in BAP1-OE hTSCs and EVTs compared to control cells. On the bottom, quantification of MMP2 activity (normalized to hTSC controls; mean ± SEM, *n*=4 independent experiments; *p<0.05, two-way ANOVA with Tukey’s multiple comparisons test). (**C**) Volcano plot of differentially expressed genes (DEGs) in BAP1-OE EVTs versus control EVTs (red: upregulated genes; blue: downregulated genes; adjusted *p*< 0.05, |log2FC| > 1). (**D**) Functional enrichment analysis of DEGs, highlighting pathways associated with upregulated and downregulated genes. Circle size represents gene count; color indicates statistical significance (-log10(p-value)). (**E**) Heatmap of FPKM-normalized expression for extracellular matrix (ECM)-related genes, showing dysregulated expression patterns in BAP1-OE EVTs (row-scaled Z-scores). OE, overexpressed.

**Supplementary Figure 4:**
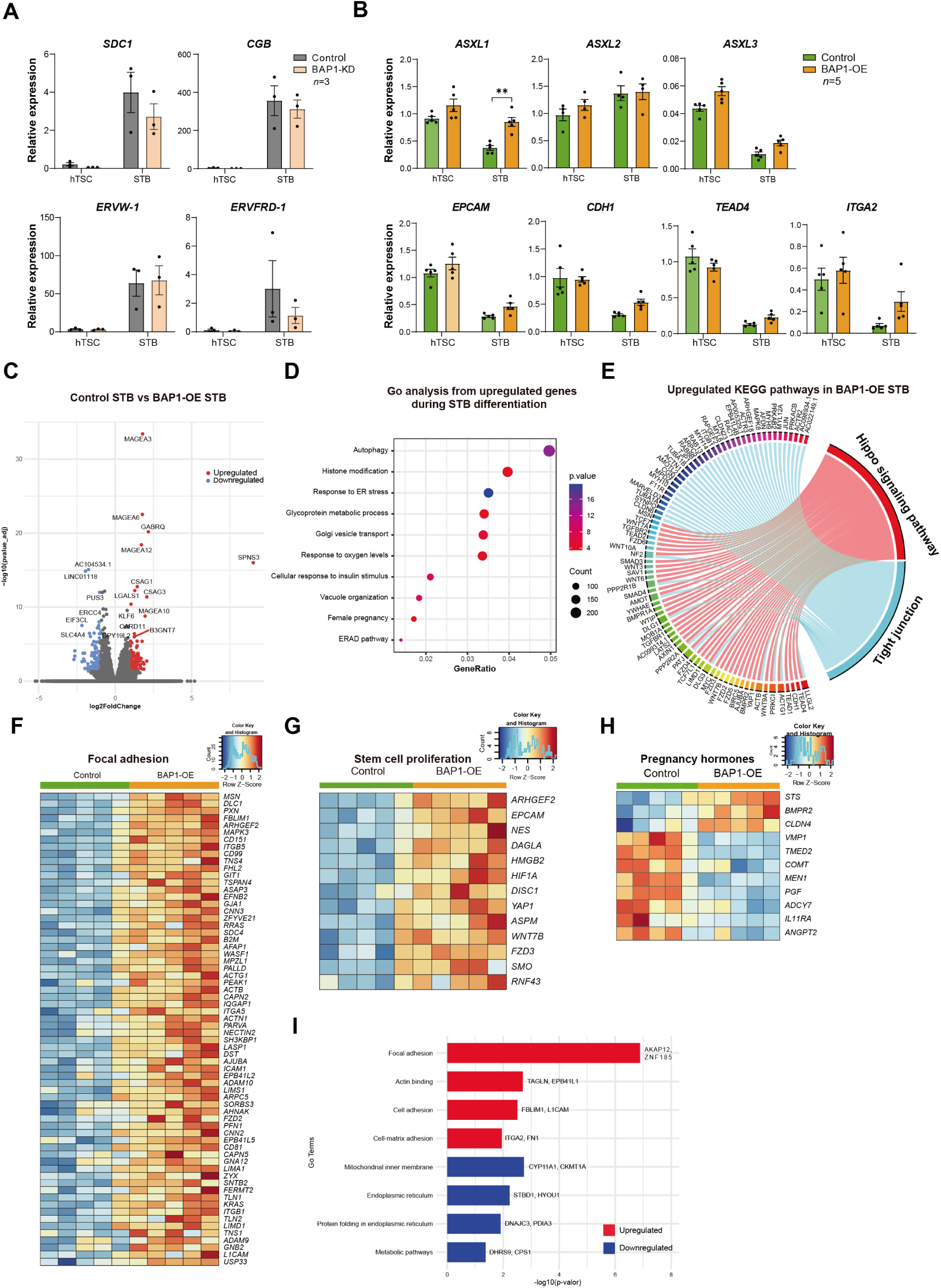
BAP1 overexpression disrupts syncytiotrophoblast differentiation. (**A**) RT-qPCR analysis of syncytiotrophoblast (STB) markers (*SDC1*, *CGB*, *ERVW-1*, *ERVFRD-1*) in BAP1-KD hTSCs and STBs compared to controls. Data are mean ± SEM*, n=*3 independent experiments, two-way ANOVA with Tukey’s multiple comparisons test. (**B**) RT-qPCR analysis of PR-DUB components (*ASXL1-3*), epithelial (*EPCAM*, *CDH1*) and stem cell (*TEAD4*, *ITGA2*) markers in BAP1-OE hTSC and STBs compared to controls (mean ± SEM of *n=*5 independent experiments; \*\**p*<0.005, two-way ANOVA with Tukey’s multiple comparisons test). (**C**) Volcano plot of significantly upregulated (red) and downregulated (blue) genes (|log2FC>1|, adjusted *p*<0.05) in BAP1-OE compared control STBs. (**D**) Gene Ontology (GO) enrichment analysis (bubble plot) of upregulated pathways upon normal STB differentiation. Circle size represents gene count; color indicates statistical significance (-log10(p-value)). (**E**) Chord diagram illustrating key upregulated pathways in BAP1-OE STBs compared to Control STBs. (**F-H**) Heatmaps of FPKM values for genes related to (**F**) focal adhesion, (**G**) stem cell proliferation, and (**H**) pregnancy hormones in BAP1-OE and control samples. Rows are z-score normalized. (**I**) Functional enrichment analysis of DEPs, highlighting pathways associated with upregulated (red) and downregulated (blue) proteins. hTSC, human trophoblast stem cell; OE, overexpression.

**Supplementary Figure 5:**
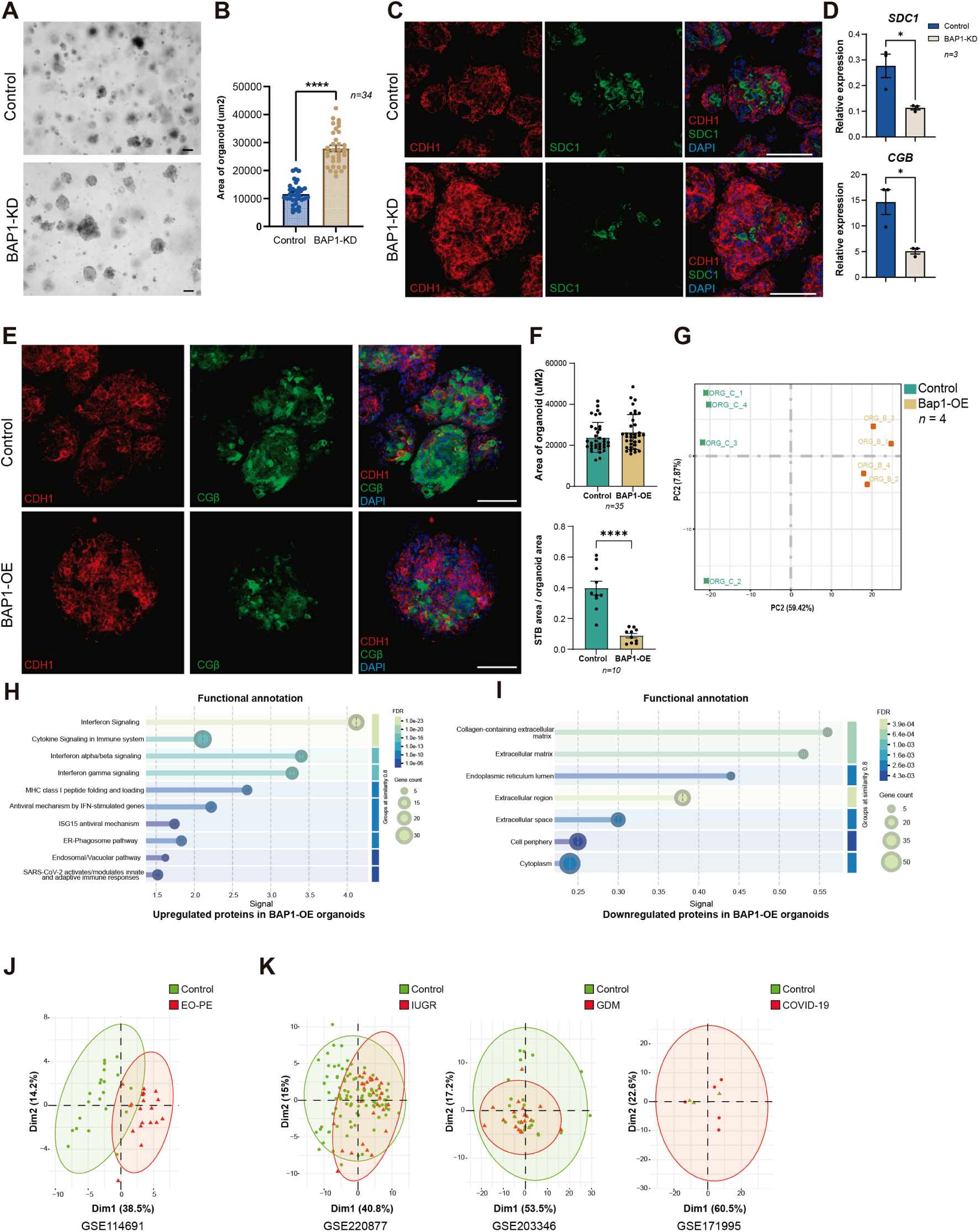
BAP1 overexpression disrupts trophoblast organoid formation and recapitulates early onset-preeclampsia features. (**A**) Brightfield images demonstrating increased organoid size in BAP1-KD compared to controls. Scale bars, 100 µm. (**B**) Quantification of total organoid area, confirming morphological expansion in BAP1-KD conditions (mean ± SEM, *n=*34 organoids/group; p****<0.0001; Student’s two-tailed t-test) (**C**) Immunofluorescence analysis of epithelial (CDH1) and syncytiotrophoblast (SDC1) markers in BAP1-KD organoids compared to controls, with DAPI nuclear counterstain. Scale bars, 100 µm. (**D**) RT-qPCR profiling of syncytiotrophoblast markers (*SDC1, CGB*) in BAP1-KD versus control organoids (mean ± SEM, *n=*3 independent experiments; *p<0.05; Student’s two-tailed t-test). (**E**) Immunofluorescence analysis of BAP1-OE and control organoids for CDH1 and CGβ. Nuclear counterstaining was with DAPI. Scale bars, 150 μm. (**F**) Quantification of organoid cross-sectional area (upper graph) shows no significant differences between groups (mean ± SEM; *n* = 35; Student’s two-tailed *t*-test). The lower graph shows the ratio of syncytiotrophoblast (STB) area to total organoid area, revealing a significant reduction in STB formation in BAP1-OE organoids (mean ± SEM; *n* = 10; **** *p* < 0.0001; Student’s two-tailed *t*-test). (**G**) Principal component analysis (PCA) of RNA-seq data showing distinct clustering of BAP1-overexpressing and control organoids, reflecting global transcriptional differences. (**H-I**) STRING functional annotation analysis of (**H**) upregulated and (**I**) downregulated pathways in BAP1-OE organoids compared to control organoids. Circle size represents gene count; color indicates False Discovery Rate (FDR). (**J-K**) Principal component analysis (PCA) plots showing separation between control (green) and disease (red) placental samples from EO-PE (GSE114691) (**J**), and IUGR (GSE220877), GDM (GSE203346), and COVID-19 (GSE171995) datasets (**K**), based on the BAP1-OE molecular signature. Axes represent principal components 1 and 2, explaining the indicated percentage of total variance. OE, overexpression.

